# A Structural Domain in the genomic RNA of SARS-CoV-2 Folds into a Compact Granular Structure without the N protein: A Single-Molecule Fluorescence Spectroscopic Investigation

**DOI:** 10.64898/2026.05.15.725307

**Authors:** Takahiro Kimura, Takuya Katayama, Shion Ishikawa, Shrutarshi Mitra, Yuuhei Yamano, Kazumitsu Onizuka, Fumi Nagatsugi, Shilpi Laha, Athi N. Naganathan, Yuji Itoh, Satoshi Takahashi

**Affiliations:** Institute of Multidisciplinary Research for Advanced Materials, Tohoku University, Sendai, Miyagi, Japan; Graduate School of Life Sciences, Tohoku University, Sendai, Miyagi, Japan; Department of Chemistry, Graduate School of Science, Tohoku University, Sendai, Miyagi, Japan; Department of Biotechnology, Bhupat & Jyoti Mehta School of Biosciences, Indian Institute of Technology Madras, Chennai 600036, India

**Keywords:** genomic RNA, RNA structure, SARS-CoV-2, N protein, genome packaging, single-molecule FRET spectroscopy

## Abstract

Severe acute respiratory syndrome coronavirus 2 (SARS-CoV-2) packages its single-stranded genomic RNA (gRNA) having about 30,000 nucleotides into virions by forming 35–40 granular ribonucleoprotein (RNP) units. Each RNP unit has a diameter of ∼15 nm. While it is generally assumed that the assembly of these RNPs is driven by the binding of the nucleocapsid (N) protein to the gRNA in the cytoplasm, the precise molecular mechanism remains to be fully elucidated. In this study, we develop an experimental strategy based on single-molecule fluorescence and fluorescence correlation spectroscopies to examine the formation of long-range base pairing within a candidate structural domain corresponding to nt 12230-12686 of the gRNA (gRNA_12k_). Our results demonstrate that the 5’ and 3’ regions of gRNA_12k_ autonomously form long-range base pairing in near-physiological buffers containing mono- and divalent cations, independently of the N protein. This domain possesses an extensive secondary structure, is compact, and can unfold and refold reversibly upon heat treatment and cooling. Notably, the addition of the N protein melts the long-range base pairs, and causes the aggregation of multiple molecules of gRNA_12k_. Based on these observations, we propose a refined mechanism for the genome assembly in SARS-CoV-2: gRNA initially forms autonomous granular structures, which are subsequently reorganized and condensed by the N protein to chaperone the assembly of the entire gRNA.

**Significance:** SARS-CoV-2 organizes its exceptionally long genomic RNA (gRNA) having about 30,000 nt into 35–40 granular ribonucleoprotein (RNP) units for viral packaging. It has been assumed that the nucleocapsid (N) protein drives the formation of the RNP granules. In this study, we challenge this prevailing view by demonstrating that a specific region of the gRNA sequence inherently encodes the information to fold into a compact, granular architecture independently of any proteins. Unexpectedly, we found that the N protein partially melts the autonomous structures, suggesting that it acts as an RNA chaperone to facilitate flexible genome assembly. Our findings redefine the interplay between viral proteins and gRNA, offering a new perspective on the mechanism of coronavirus replication.

## Introduction

The packaging of genomic RNA (gRNA) into lipid membranes of the endoplasmic reticulum–Golgi intermediate compartment (ERGIC) is a fundamental stage in the replication of severe acute respiratory syndrome coronavirus 2 (SARS-CoV-2) (1–4). As a positive-sense, single-stranded RNA virus with a huge genome having about 30,000 nucleotides (nt), SARS-CoV-2 is considered to organize its genetic material via the nucleocapsid (N) protein (5–11). This association results in the formation of 35–40 granular ribonucleoprotein (RNP) units that are densely packed within the viral envelope (12, 13). Each RNP complex having a diameter of ∼15 nm is thought to consist of ∼12 molecules of the N protein and ∼800 nt region of the gRNA (12, 14). While current models, including that hypothesized by ourselves suggest that the assembly of the RNP granules is driven by the N protein-gRNA interactions in the cytosol (9, 12–16), the precise biophysical principles of this process remain elusive. Here, we challenge this view by demonstrating that a region of the gRNA sequence inherently encodes the information required to fold into granular architectures, independent of protein scaffolds.

Various methods suggested the presence of granular RNP structures inside of SARS-CoV-2 virus. The cryo-electron microscopy (cryo-EM) provided pivotal insights into the granular architecture of these RNPs (12, 13). In some analyses, each RNP particle was suggested to be organized as a “crown-like” assembly of 10–12 molecules of the N protein, which is encapsulated by gRNA (12–14). These observations are complemented by *in vitro* studies showing that purified N protein and the stem loop region of gRNA spontaneously assemble into granules having a dimension similar to that of the RNPs (14, 17, 18). Concurrently, viral RNA *in situ* conformation sequencing (vRIC-seq) has revealed that the gRNA forms an extensive network of secondary structures, predominantly stem-loops ranging from 40 to 80 nt (19). In addition, approximately 49 long-range base-pairing interactions spanning the median distance of ∼630 nt have been identified across the genome (19). This numerical correspondence with the estimated 35–40 RNP granules suggests a hypothesis: the “domains” of gRNA, defined as regions of gRNA that are enclosed by the long range base pairing detected in the vRIC-seq analysis, corresponds to the RNP “granules”, defined as the spherical units having a diameter of ∼15 nm, and the N protein may recognize the pre-existing stem-loops in the gRNA, thereby facilitating the folding of the domains and stabilizing the granules through the formation of long-range base pairs. However, direct experimental evidence for such a coordinated assembly process remains absent.

Intriguingly, entirely different biophysical phenomena have been reported regarding the N protein-RNA association. First, when mixed with relatively short RNA fragments, the N protein frequently undergoes liquid-liquid phase separation, forming droplets that can exceed the dimensions of the virion itself (18, 20–23). The physical properties of these condensates appear to be RNA-dependent: while single-stranded RNA fragments tend to form low-viscosity droplets, sequences containing stem-loops result in high-viscosity, amorphous aggregates (24). Second, the N protein works as an RNA chaperone and promotes both melting and annealing of RNA duplexes (25–29). Together, these observations rather suggest that the association between the N protein and gRNA is non-specific.

The current disparity between these structural and biophysical observations underscores the need for experimental methodologies capable of resolving the molecular-level assembly process. To bridge this gap, we employed single-molecule fluorescence resonance energy transfer (sm-FRET) spectroscopy (30–32) and fluorescence correlation spectroscopy (FCS) (33, 34) to analyze the conformational dynamics of gRNA_12k_ corresponding to nt 12230-12686 of gRNA. By labeling the 5′ and 3′ ends of gRNA_12k_ with fluorescent dyes, we were able to monitor the inter-terminal distance as a proxy for the long-range base pair formation. Contrary to our initial hypothesis, we found that gRNA_12k_ can form the long-range base pairs independently of the N protein. Furthermore, we demonstrated that the association of the N protein to gRNA_12k_ impedes these long-range base pairs. Based on these findings, we propose a new model that redefines the interplay between the N protein and the gRNA sequence in SARS-CoV-2 assembly.

## Results

### Selection, preparation and labeling of the candidate granular domain of gRNA

The vRIC-seq analysis showed that gRNA of SARS-CoV-2 in virions harbors multiple long-range base pairs, which likely correspond to compact topological domains of gRNA (Fig 1A) (19). A total of 49 of long-range base pairs were identified, with a length of base-paired region of less than 40 bp and a median separation of ∼630 nt. Thus, we hypothesized that the sequences separated by several hundred nucleotides interact with each other to form topological domains in the gRNA. While the approaches such as vRIC-seq can infer the secondary structures, they cannot directly observe tertiary structures. To determine how or under what conditions the long-range base pairs form, we adapted sm-FRET spectroscopy by labeling donor and acceptor fluorophores at the two terminal nucleotides of a representative topological domain (35–37).

**Figure 1.**
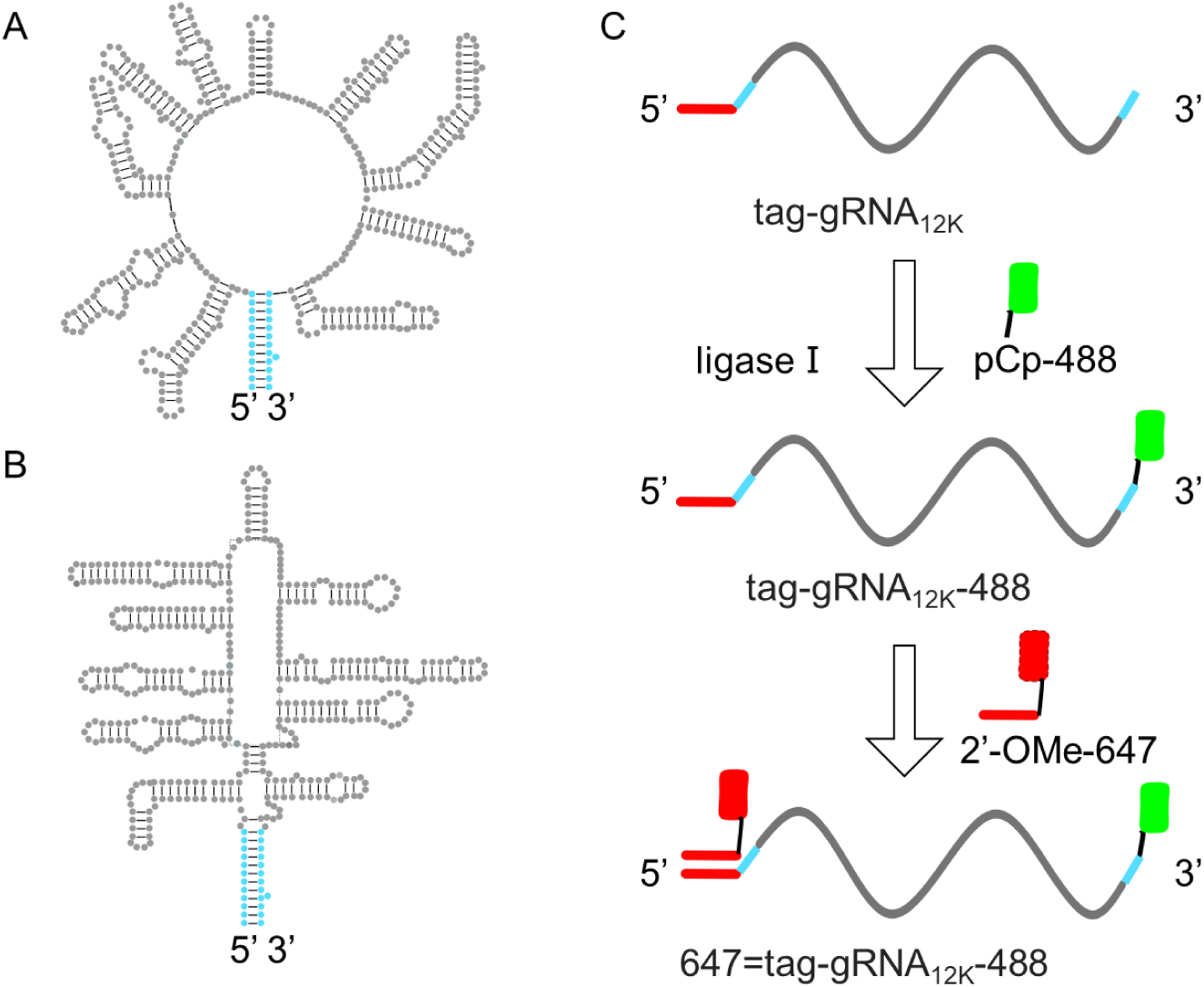
Proposed secondary structures of gRNA_12k_ and its labeling strategy. (A) and (B) Secondary structure maps of gRNA_12k_ deduced using the vRIC-seq (A) and SHAPE-MaP (B) methods. (C) The 494-nt RNA sample corresponding to the 5′-end 22-nt tag followed by the 12230-12686 region of gRNA, tag-gRNA_12k_, was prepared by *in vitro* transcription. The 3′ terminus was ligated with pCp-Alexa488 using ligase I (tag-gRNA_12k_–488). A 22 nt 2′-*O*-methylribonucleotide with Alexa647 at the 5′ terminus, whose sequence was complementary to the 5′-end 22-nt tag (2′-OMe-647), was hybridized (647=tag-gRNA_12k_–488). The sequence information and the gel analysis of the labeling and purification process were presented in Supporting Table S1 and Supporting Fig. S1, respectively.

We chose the 457-nt region spanning nt 12230 to 12686 of gRNA, gRNA_12k,_ as a target of this investigation. The vRIC-seq analysis showed that two terminal sequences of gRNA_12k_, nt 12230-12241 and nt 12674-12686, formed base pairs with a high confidence (Fig 1A) (19). The same long-range base pair and the same number of stem-loops between them were also determined by using selective 2′-hydroxyl acylation analyzed by primer extension and mutational profiling (SHAPE-MaP) (Fig 1B) (38). This is the only domain for which both vRIC-seq and SHAPE-MaP analyses suggest the same long-range base pairs. Given the consistency of the results, we chose gRNA_12k_ as the model domain for detailed spectroscopic analyses.

We labeled the 5′ and 3′ ends of gRNA_12k_ with Alexa647 and Alexa488, respectively. A 494-nt RNA sample corresponding to gRNA_12k_ with a 31-nt tag extension at the 5′ end and an additional 6-nt extension (tag-gRNA_12k_) was synthesized by using *in vitro* transcription (Fig 1C, Supporting Table S1). pCp-Alexa488 was ligated to the 3′ terminus of the sample (tag-gRNA_12k_–488) (36). A 22-base 2′-*O*-methyl ribonucleotide having a sequence complementary to the 5′ tag and Alexa647 at its 5′-terminus (2′-OMe-647) was hybridized to the 5′ tag of tag-gRNA_12k_–488 (647=tag-gRNA_12k_–488) (39, 40). Finally, the samples were purified using an agarose gel (Supporting Fig S1). The details of the sample preparation are discussed in the Supporting Text.

### Proximity of the 3’ and 5’ termini of 647=tag-gRNA_12k_–488 in the presence of mono- and divalent cations

When the distance between the donor and acceptor fluorophores falls within the range of 1–10 nm, the resonance energy transfer of the excited singlet state occurs from the donor to the acceptor, whose transfer efficiency (*E*), determined using the sm-FRET spectroscopy can be used to estimate distance between the fluorophores (30–32). If the long-range base pairing occurs in 647=tag-gRNA_12k_–488, a high efficiency population would be expected. We first conducted the sm-FRET measurements of 50-pM 647=tag-gRNA_12k_–488 in a near physiological buffer containing 150 mM NaCl and 1 mM of MgCl_2_ (41). Figure 2A shows a two-dimensional (2D) histogram whose horizonal and vertical axis correspond to *E* and the stoichiometry (*S*) of each burst, respectively, determined by the alternating-laser excitation (ALEX) detection method of the sm-FRET spectroscopy (42, 43). The *S* value describes the ratio of photon numbers detected during the donor excitation and during the acceptor excitation (42, 43). We eliminated bursts having the *S* values near 0 or 1, since they correspond to the acceptor only or donor only species, respectively. The 2D plot indicates that all the selected bursts possess the *S* values from 0.3 to 0.7, demonstrating that they originate from species having the both fluorophores. The FRET efficiency distribution possessed peaks at 0.3, 0.6 and 0.8, in addition to the peak at 0, suggesting that a significant fraction of the sample possesses conformations in which the 5′ and 3′ termini are in the proximity.

**Figure 2.**
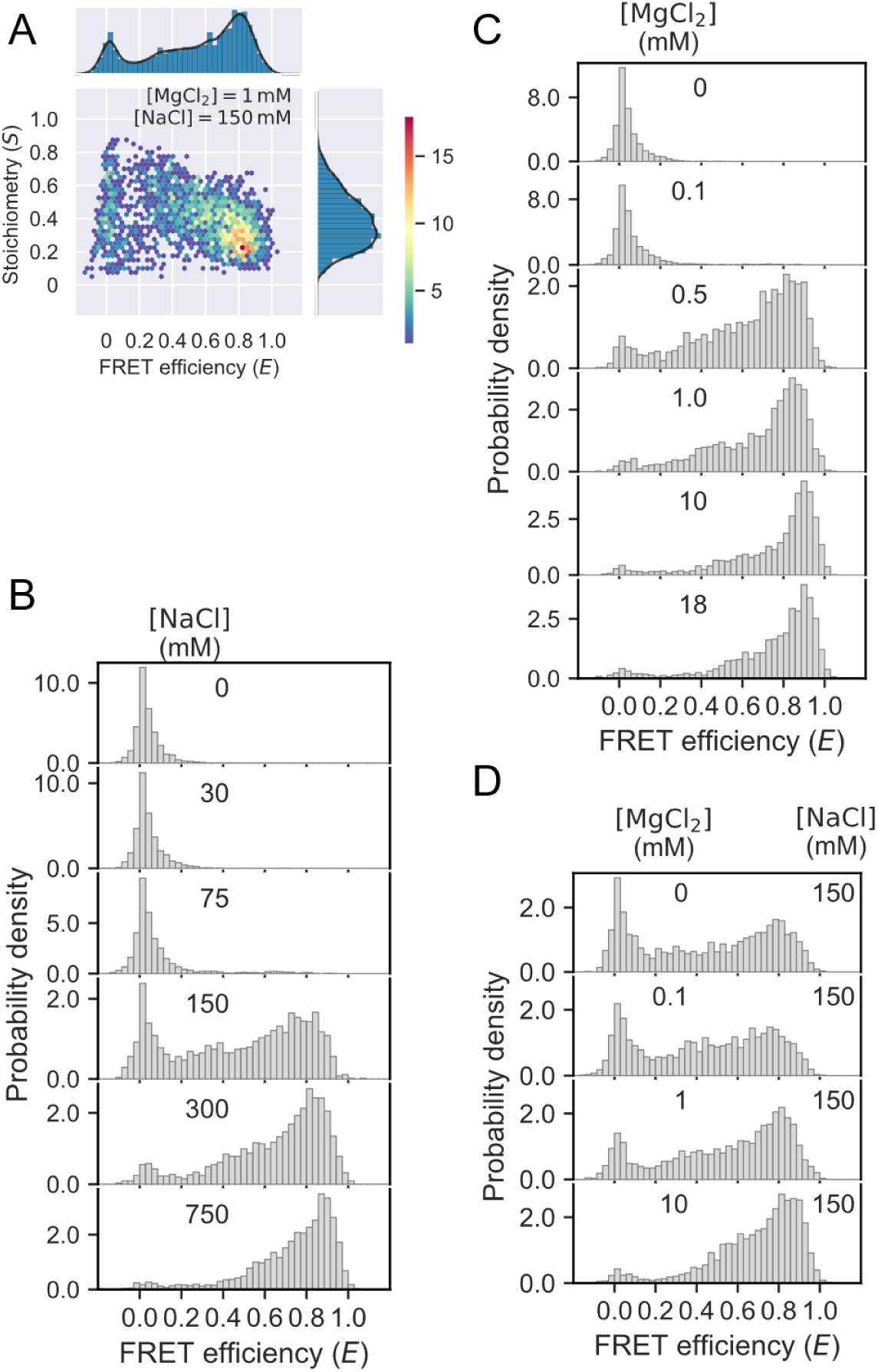
The sm-FRET spectroscopic measurements for 50-pM 647=tag-gRNA_12k_–488 at various concentrations of sodium and magnesium ions. (A) A two-dimensional *E*-*S* plot for the sample obtained in the presence of NaCl and MgCl_2_ at 150 mM and 1 mM, respectively. (B) Changes in the sm-FRET efficiency at different concentrations of NaCl in the absence of MgCl_2_. (C) Changes in the FRET efficiency at different concentrations of MgCl_2_ in the absence of NaCl. (D) Changes in the FRET efficiency at different concentrations of MgCl_2_ in the presence of 150 mM NaCl. More than 1700 bursts were used to depict each of the distributions presented in panels B–D. The burst width distributions corresponding to the efficiency distributions were shown in Supporting Fig. S2.

Since RNA structures depend strongly on ionic strength, we next investigated the sm-FRET efficiency distributions at different concentrations of Na^+^ (Fig. 2B and Supporting Fig. S2). At concentrations of Na^+^ less than 75 mM, only the *E* = 0 species was detected. At 150 mM, the bursts having the efficiencies at ∼0.8 increased. At the highest concentration examined (750 mM), the ∼0.85 species became the majority. Assuming the free rotation of Alexa488 and Alexa647 and a fixed distance between them, and using the Förster distance (*R*_0_) of 55 Å for the Alexa488−Alexa647 pair (44), the species with the efficiencies of less than 0.05, 0.3, 0.6 and 0.85 correspond to the donor and acceptor distances of more than 9.0 nm, 6.3, 5.1 and 4.1 nm, respectively. These results demonstrate that the screening of the electrostatic repulsion with Na^+^ triggers the structural transition of the sample, resulting in multiple conformations with relatively short distances between the 5′ and 3′ termini.

We next investigated the effect of the divalent cation, Mg^2+^, on the sm-FRET efficiency distributions (Fig. 2C and Supporting Fig. S2). Mg^2+^ is known to stabilize RNA structures at much lower concentrations than monovalent cations (37, 45–48) . At 0.1 mM Mg^2+^, only the *E* = 0 species was detected. At 0.5-mM Mg^2+^, the bursts having higher FRET efficiencies increased. At concentration higher than 0.5 mM, the *E* = ∼0.9 species became dominant. With the constant Na^+^ concentration of 150 mM, the sm-FRET distribution was not strongly dependent on the Mg^2+^ concentration up to 1 mM (Fig. 2D and Supporting Fig. S2). However, the species with higher efficiencies increase at higher concentrations of Mg^2+^ (see discussion below). These data indicate that the divalent Mg^2+^ stabilizes the compaction of gRNA_12k_ more efficiently than the monovalent Na^+^, confirming that the non-specific screening of the electrostatic repulsion drives the structural transition.

### Formation of the long-range base pairs in 647=tag-gRNA_12k_–488

The proximity of the 3′ and 5′ termini detected for 647=tag-gRNA_12k_–488 suggests that the domain formed the long-range base pairing. However, it is still possible that the proximity reflects non-specific chain compaction due to the screening of the electrostatic repulsion. To verify the presence of the long-range base pairing, we adopted the antisense strategy proposed previously (35) and performed the sm-FRET measurements for 80-pM 647=tag-gRNA_12k_–488 in the presence of antisense oligonucleotides (ASO1). We prepared 13-nt 2′-*O*-methylribonucleotides (ASO1) and 12-nt ASO2, whose sequences were complementary to the 3′ base-pairing region (nt 12674-12686) and to the 5′ region (nt 12230-12241) of gRNA_12k_, respectively (Figs. 3A and 3C), and performed the sm-FRET measurements of 647=tag-gRNA_12k_–488 in the presence of ASO1 and ASO2 (See Supporting Text and Supporting Table S1 for the detailed protocol and sequence information, respectively). In addition, we conducted control measurements in the presence of a non-specific 13-nt poly-adenylate 2′-*O*-methylribonucleotides (rA_13_). As shown in Fig. 3B and 3D, the peaks at *E* = ∼0.4 and ∼0.8, which were observed in the absence of ASOs, diminished with the increase of the ASO1 or ASO2 concentrations. The effect was not detected for rA_13_, clearly indicating that the middle and high efficiency populations are associated with the base pairing between the 5′ and 3′ regions. These results demonstrate that the long-range base pairing of the sample is formed in the presence of mono- and divalent cations.

**Figure 3.**
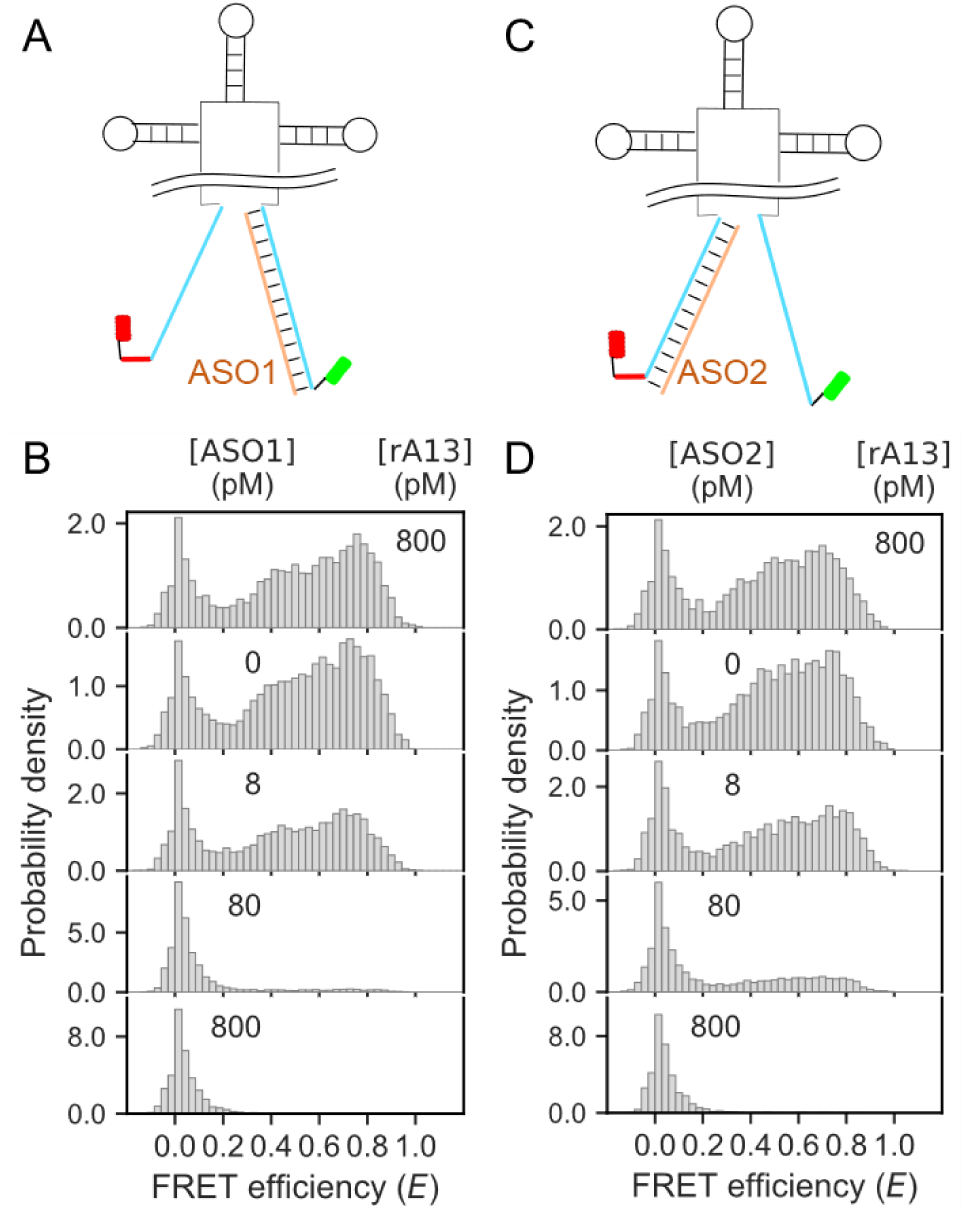
The sm-FRET spectroscopic measurements for 80-pM 647=tag-gRNA_12k_–488 in the presence of ASOs. (A and C) A schematic illustration of ASO1 (A) and ASO2 (C) binding to the sequences constituting the long-range base pairing, thereby disrupting FRET. (B and D) The sm-FRET efficiency distributions obtained at different concentrations of ASO1 (B) or ASO2 (D) were presented. As a control, the sample incubated with rA_13_ without ASOs were presented at the top. Detailed procedure of the sample preparation was discussed in Supporting Text and Supporting Fig. S3. The observation buffer contained 150 mM NaCl and no MgCl_2_.

### gRNA_12k_ is compact, rich in secondary structures and shows a reversible thermal melting

To determine the compactness of gRNA_12k_ by the FCS measurements at the 488-nm excitation, we hybridized a non-labeled 2′-*O*-methylated oligonucleotide to the 5′-tag of the singly labeled tag-gRNA_12k_–488 (2’-OMe=tag-gRNA_12k_–488). The FCS data of the sample obtained in the solution containing both 150 mM Na^+^ and 1 mM Mg^2+^ showed that *R*_H_ was 7.1 ± 1.3 nm (Supporting Fig. S4 and Supporting Tables S2 and S3). In the absence of the cations, *R*_H_ expanded to 10.1 ± 1.0 nm. An expansion of ssRNA in the absence of salt has been reported earlier (48, 49). The compaction of gRNA_12k_ in the presence of the cations was similarly reflected in the burst width distributions, showing the narrowing of the width upon the increase of NaCl concentration (Supporting Fig. S2). To investigate the secondary structure content, we performed the circular dichroism (CD) measurements of tag-gRNA_12k_. The CD spectrum for tag-gRNA_12k_, dissolved in 20 mM tris buffer at pH 7.5, displays a strong positive ellipticity at 263 nm (Fig. 4A), resembling that of stem loop RNAs. We compared the peak ellipticity of tag-gRNA_12k_ to that of SL4, a fragment of gRNA corresponding to nt 78 to 127 and forming the 4^th^ stem loop (15). Assuming the formation of 17 base pairs in SL4, ∼125 base pairs might be formed in tag-gRNA_12k_, which is roughly consistent with the number of the base pairs in gRNA_12k_ deduced based on the vRIC-seq analysis (19). The CD spectra showed slight shifts in the peak wavelengths in the presence of varying concentrations of Na^+^ and Mg^2+^; however, the ellipticities remained mostly constant. This implies that the stem loop content in tag-gRNA_12k_ does not change significantly with the addition of the cations (Fig. 4A and Supporting Fig. S5A).

**Figure 4.**
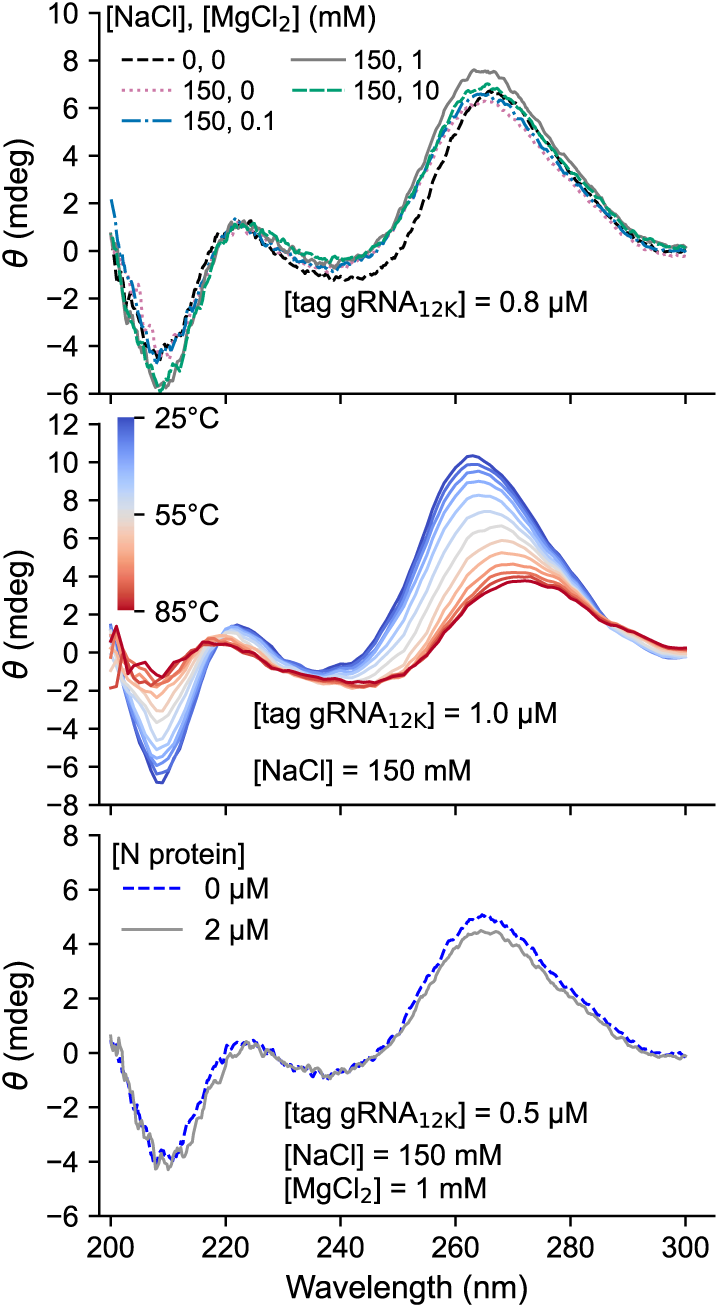
CD spectra of tag-gRNA_12k_. (A) CD spectra of 0.8 µM gRNA_12k_ in the presence of various concentrations of NaCl and MgCl_2_. The buffer contained 20 mM Tris-HCl at pH 7.5. Spectra obtained at different concentrations of NaCl were reported in Supporting Fig. S5A. (B) CD spectra of tag-gRNA_12k_ in the presence of 150 mM Na^+^ measured at temperatures from 25 to 85°C. (C) CD spectra of 0.5 µM tag-gRNA_12k_ at 150 mM Na^+^ and 1 mM of Mg^2+^ in the absence and presence of the N protein at 2 μM. The CD spectra of the N protein obtained at the respective concentration were subtracted.

To examine the stability of the stem loops in gRNA_12k_, we examined the thermal denaturation of the unlabeled sample (tag-gRNA_12k_) by observing its CD spectra at every 5°C from 25°C to 85°C (Fig. 4B). As the temperature increased, the spectra showed a decrease in ellipticity, demonstrating the gradual melting of the secondary structures in the range from 45°C to 70°C. The sigmoidal transition, as well as the presence of the isoelliptic points at 218, 234, and 285 nm, suggests that the melting occurs relatively cooperatively. In addition, we confirmed that the CD spectrum of the sample, once heated up to 85°C and cooled to 25°C, overlapped perfectly with that of the initial sample (Supporting Fig. S5B). To further investigate the reversibility, 647=tag-gRNA_12k_–488 was heat denatured for 5 min, incubated at 37°C for 5 min, and examined using the sm-FRET spectroscopy at room temperature. We initially heat denatured the sample at 85°C. However, the number of bursts observed for the sample after the cooling decreased significantly, suggesting that 2′-OMe-647 disassociated from the sample (data not shown). On the other hand, the number of bursts for 647=tag-gRNA_12k_–488 initially heat incubated at 75°C for 5 min and measured after cooling to room temperature was similar to that observed for the samples without the heat treatment. Furthermore, the sm-FRET efficiency distributions were identical to those without heat denaturation, including the Na^+^ and Mg^2+^ dependencies (Supporting Fig. S6). These data suggest that the long-range base pair formation for gRNA_12k_ is a reversible process, and that gRNA_12k_ constitutes a cooperative folding unit.

### The long-range interaction in 647=tag-gRNA_12k_–488 was abolished by the addition of the N protein

To examine how the N protein affects the folding of gRNA_12k_, we performed sm-FRET measurements of 100-pM 647=tag-gRNA_12k_–488 in the presence of the N protein and 150 mM Na^+^ (Fig. 5A and Supporting Fig. S7). Quite unexpectedly, the FRET efficiency distributions in the presence of more than 10 nM of the N protein almost completely abolished the FRET efficiency peak at ∼0.8 and increased the peak at *E* = 0. We confirmed that the peak at *E* = 0 originated from the species containing both donor and acceptor fluorophores (Supporting Fig. S7). Even in the presence of 10-mM Mg^2+^ and 150-mM Na^+^, the addition of the N protein at 100 nM and 1000 nM diminished the population having the high efficiencies (Supporting Fig. S8). These results suggest that the N protein interferes with the long-range base pair formation of 647=tag-gRNA_12k_–488 rather than facilitating it. To examine the effect on the secondary structures, we measured the CD spectra of gRNA_12k_ in the presence of the N protein (Fig. 4C). The ellipticities of the sample at 260 nm decreased for ca. 10%, showing that a part of the secondary structures was melted by the addition of the N protein. We conclude that the N protein melts the long-range base paring of gRNA_12k_ as well as some of the stem loops.

**Figure 5.**
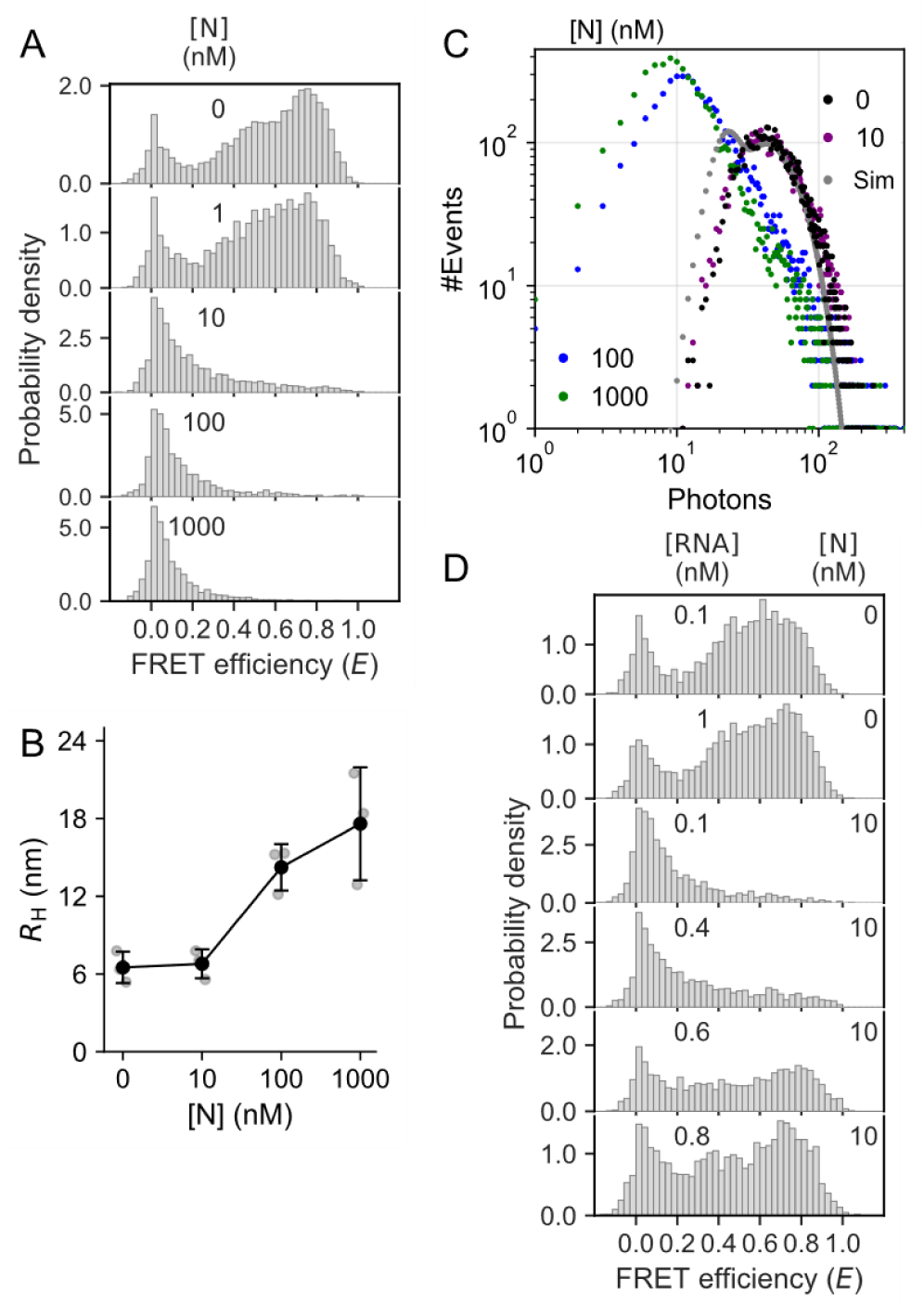
Structural transitions and aggregation of gRNA_12k_ triggered by the association of the N protein. (A) The FRET efficiency distributions for 0.1 nM 647=tag-gRNA_12k_–488 in the varying concentrations of the N protein obtained at 150 mM Na^+^ in the absence of Mg^2+^. The corresponding two-dimensional *E*-*S* plots and burst width distributions were shown in Supporting Fig. S7. Those in the presence of Mg^2+^ were shown in Supporting Fig. S8. (B) Changes in *R*_H_ in the presence of the N protein obtained by analyzing the three independent FCS data for 2’-OMe=gRNA_12k_-488 excited at 484 nm. While we could observe the significant increase in *R*_H_ at the higher concentrations of the N protein, the observed values changed significantly in the different measurements. As we discussed in Supporting Text, this might be caused by the fluctuation of the sample conditions such as the adsorption of aggregates onto the surface of the glass-based dish. The detailed fitting parameters are presented in Supporting Table S4. The averaged *R*_H_ values are presented in Supporting Table S5. (C) The fluorescence photon intensity histograms of 2-nM 488=tag-gRNA_12k_–647 in the presence of 1 mM Mg^2+^ and 150 mM Na^+^. The data obtained in the presence of 0 nM, 10 nM, 100 nM and 1000 nM of the N protein were shown in black, purple, blue and green, respectively. Gray dots represent the simulated histogram depicted by assuming a single emitting component based on the brightness and number of the RNA molecules estimated for the data in the absence of the N protein as explained in Supporting Text. (D) The FRET efficiency distributions for 0.1 nM 647=tag-gRNA_12k_–488 in the varying concentration of the labeled and non-labeled RNAs in the presence of 10 nM N protein. The total concentration of the labeled and non-labeled RNA samples was presented in each panel. As controls, the data in the absence of the N protein at 0.1 nM of 647=tag-gRNA_12k_–488, and in the absence of the N protein at 1 nM of the total RNA sample were presented. The burst width distribution plots of the same data set were presented in Supporting Fig S9.

While gRNA_12k_ remained monomeric in the absence of the N protein, the addition of the N protein caused its aggregation. To examine *R*_H_, we initially performed FCS measurements on 2 nM 2’-OMe=tag-gRNA_12k_–488 at an excitation wavelength of 484 nm (Fig. 5B). At the N protein concentrations up to 10 nM, the sample in the presence of 1-mM Mg^2+^ and 150-mM Na^+^ remained monomeric having *R*_H_ of ∼6.8 nm. While *R*_H_ increased more than twice at higher N protein concentrations, the values might represent minor species since the Alexa488 fluorescence was strongly quenched by the N protein (15). We subsequently prepared 2-nM 488=tag-gRNA_12k_–647 and monitored its fluorescence fluctuations under 642-nm excitation. Since Alexa647 fluorescence is less susceptible to quenching by N protein binding, we were able to detect numerous “spike-like” intensity fluctuations, apparently corresponding to aggregated species. However, the spikes precluded standard FCS analysis. Instead, we conducted the “brightness and number” analysis of the fluorescence intensity fluctuations detected at 10-ms time interval (50). For the N protein concentrations of 0 nM and 10 nM (shown as black and purple points in Fig. 5C, respectively), the histograms of the fluctuation data exhibited single peaks at ∼50 photons. The distribution was semi-quantitatively reproduced by assuming that a free fluorophore (∼35% of the total photon counts) contributing to a constant background and a single emitting particle having the brightness (21.8) and the number (1.6) determined from the statistics of the fluctuation (Supporting Text). This brightness can serve as a standard for monomeric fluorophore. The rapid decay of the histogram at higher photon counts is typical of single emitter obeying a Poisson distribution. At higher N protein concentrations, the peak intensity of the histogram shifted to lower photon counts, indicating that the number of fluorescent particles including free fluorophores was reduced. The averaged brightness and number of the RNA molecules were determined to be 79.7 and 0.173, respectively, assuming the presence of ∼30% free fluorophore. The brightness suggests that ∼4 (79.7/21.8) fluorophores are present in a single particle. Since the labeling efficiency was ∼60%, ∼6 RNA molecules were aggregated. The histogram at higher photon counts decreased at a slower rate than that observed in the absence of the N protein, showing the presence of multiple oligomeric states. Finally, the sm-FRET measurements of 0.1 nM 647=gRNA_12k_–488 showed the significant expansion of the burst width in the presence of the N protein (Supporting Figs S7 and S8). These results demonstrate that the N protein aggregates the multiple molecules of gRNA_12k_.

To determine the stoichiometric ratio of the N protein relative to gRNA required for the melting of the long-range base pairs, we added the non-labeled tag-gRNA_12k_ to change the total RNA concentration while keeping the concentration of 647=tag-gRNA_12k_–488 constant at 100 pM (Fig. 5D and Supporting Fig. S9). Without the addition of the non-labeled RNA, the *E* = 0 species is dominant in the presence of 10 nM N protein. However, the addition of the non-labeled RNA at the ratio of the N protein relative to the total RNA ([N]/[RNA]_total_) of 25, the *E* = ∼0.8 species started to emerge. At the [N]/[RNA]_total_ ratio of 16, the *E* = ∼0.8 species became dominant, suggesting that the number of the N protein attached to RNA was not sufficient to melt the long-range base pairs. Note that the burst width distributions obtained in at the lower [N]/[RNA]_total_ ratio was different from that in the absence of the N protein (Supporting Fig S9), showing that the N protein was still associated with 647=tag-gRNA_12k_–488. In line with this observation, we confirmed that the aggregation of the RNA samples did not occur at the [N]/[RNA]_total_ ratio less than 17. These data indicate that melting of the long-range interaction in gRNA_12k_ and the aggregation of gRNA_12k_ require more than ∼20 molecules of the N protein.

## Discussion

The structural transitions of gRNA_12k_ characterized in this investigation can be summarized as follows (Fig. 6). The sample in the absence of the N protein in near physiological buffer (41, 51), mimicking the environment of gRNA just transported from the double membrane vesicle to cytosol (16, 52), autonomously forms the granular structures in the presence of mono- and divalent cations. The domain possesses significant secondary structures, is compact with *R*_H_ of 6.5 ± 1.2 nm and has its 5′ and 3′ regions base paired each other. The addition of the N protein more than ∼20 stoichiometric amount relative to that of gRNA_12k_, which was 10 nM N protein in the sm-FRET measurements and 100 nM in the FCS and fluorescence fluctuation measurements, disrupts the long-range base pairs and caused the aggregation. The abundant concentration of the N protein, mimicking the environment of gRNA before the encapsulation by ERGIC membrane, promoted the assembly of the granules (16, 53, 54). The current finding revealed important properties of gRNA essential for the genome packaging in SARS-CoV-2.

**Figure 6.**
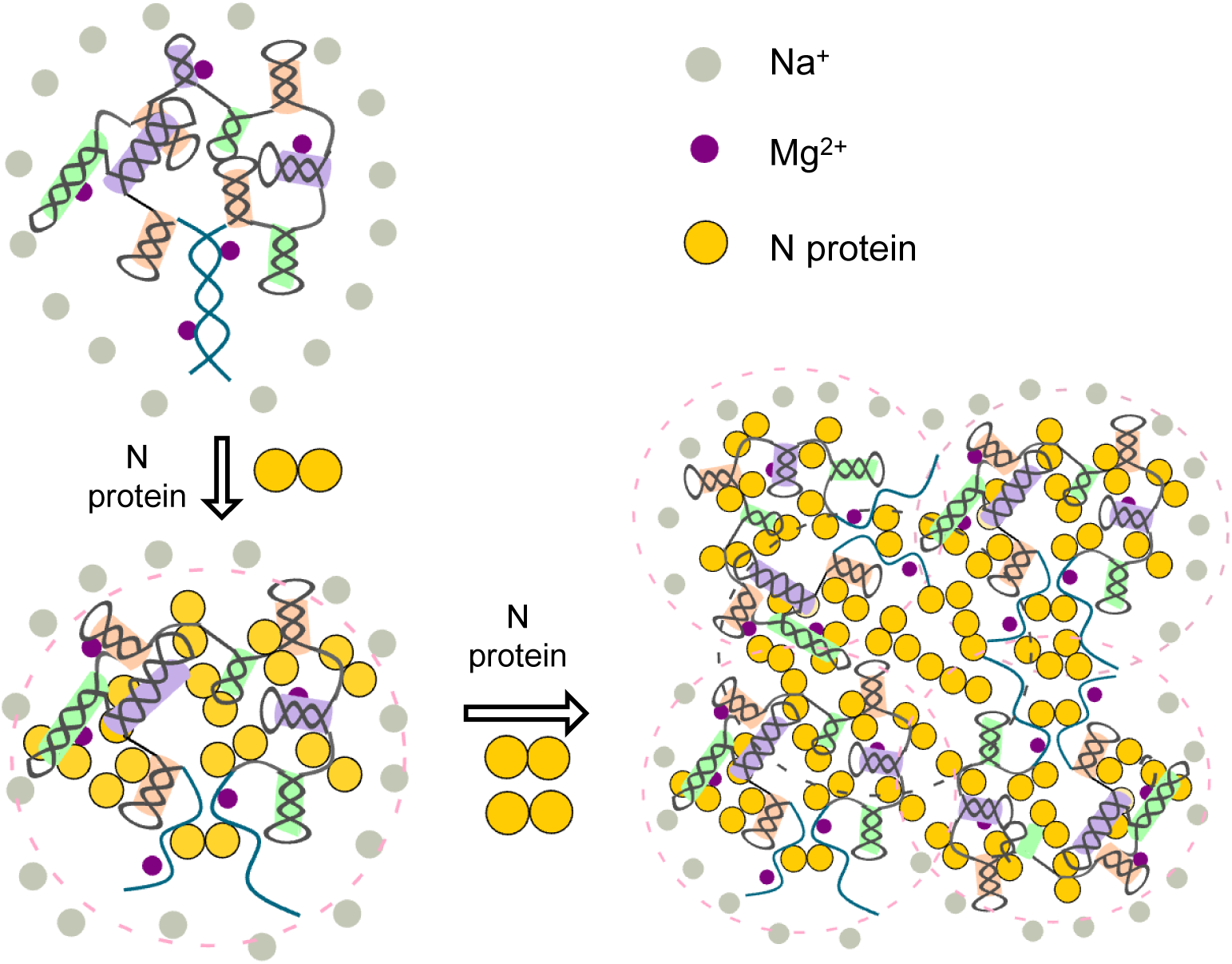
A refined model for the granular particle formation in gRNA_12k_ of SARS-CoV-2. The long-range base pairs in gRNA_12k_ are formed under near-physiological concentration of cations. The association of more than 20 molecules of the N protein disassociated the long-range base pairs and caused the aggregation of gRNA_12k_.

The existence of long-range base pairing in the gRNA of SARS-CoV-2 has been controversial. The multiple long-range interactions in the gRNA of the virus, including those of gRNA_12k_, were detected by using the vRIC-seq method (19). Further evidence of the long-range base pairing was obtained through the SHAPE-MaP analysis of the viral RNAs in host cells infected with SARS-CoV-2 (38). In contrast, the iSHAPE-MaP and SPLASH methods failed to detect the long-range base pairing in gRNA_12k_ (55, 56). These discrepancies may be due to the ensemble-averaged nature of the results based on the RNA-seq methods. The current sm-FRET measurements revealed the coexistence of several conformations (Fig. 2), among which the high FRET state corresponds to the conformations having the long-range base pairing (Fig. 3), consistent with the observations from the vRIC-seq and SHAPE-MaP methods (19, 38). In contrast, the low-FRET population might represent conformations corresponding to those detected by the iSHAPE-MaP and SPLASH methods. In addition, the N protein disrupted the long-range base pairing (Fig. 5). In vRIC-seq experiments, the gRNA in viral particles was exposed to buffers containing high concentrations of MgCl_2_ (19). Therefore, the conditions during vRIC-seq might have stabilized the long-range base paring even in the presence of the N protein. We suggest that gRNA_12k_ adopts a dynamic equilibrium between the conformations with and without the long-range base pairing, which might explain the discrepancies among the previous RNA-seq–based studies.

The structure of gRNA_12k_ is modulated by the electrostatic screening. Consistent with the reported role of Na^+^ in stabilizing RNA secondary structures by reducing electrostatic repulsion (57, 58), the high-FRET population increased with increasing Na^+^ concentration but only at the concentrations more than typical physiological ionic strength (> 150 mM) (Fig. 2B) (41). In contrast, the long-range base pairing was formed at the typical cellular concentrations of Mg^2+^ (∼1 mM) (Figs 2A, C and D) (51), consistent with the reports that the divalent cations stabilize RNA structures more efficiently than monovalent ions through both diffuse electrostatic interactions and specific coordination with the phosphate backbone (46, 59, 60). Interestingly, the effect of 150 mM Na⁺ on the pattern of the sm-FRET distribution takes precedence over that of 1 mM Mg²⁺ alone when both ions are present simultaneously. A similar observation has been reported regarding the stability of RNA helices (45), suggesting that the long-range base pairing for gRNA_12k_ occurs in the mechanism that stabilizes canonical RNA secondary structures.

We suggest that the role of the N protein in the genome packaging is not the formation of the RNP granules but is the chaperoning of the flexible assembly of the RNA granules. In general, viral ssRNAs are considered to adopt compact secondary structures for the packaging (61, 62). While we initially hypothesized that the formation of the granules occurs upon the association of the N protein, the addition of the N protein disrupted, rather than stabilized, the long-range base pairing of gRNA_12k_ and caused its aggregation. The results contradict our initial hypothesis but are consistent with the proposal that the N protein works as an RNA chaperone by partially melting and forming RNA duplexes (26–28, 63). Similar chaperone activities, characterized by the melting and forming RNA duplexes, have been documented for nucleocapsid proteins in other viruses, such as the NC protein of HIV-1 and the NP of influenza A virus (64–69). In the context of SARS-CoV-2, this chaperone activity may be essential for remodeling the autonomous, relatively rigid granular structures of the gRNA into a more flexible state. The N protein forms liquid-like droplets when mixed with short RNA fragments as well as relatively long RNA fragments (24, 70–74). A liquid like property implies that the N protein may facilitate fluidic RNA-RNA interaction, which would facilitate the high-density packaging of the massive genome within the limited internal volume of the virion.

In summary, we established the fluorescence-based experimental strategy for the structural characterization of long RNA chains under various conditions using samples preparable by *in vitro* transcription. Based on the current and previous investigations, we proposed a new mechanism of the genome packaging in SARS-CoV-2. Further research is necessary to understand the molecular principles that determine the viral life cycle. First, we must investigate other domains of gRNA. Second, alternative labeling strategies at the internal nucleotide of long RNAs are necessary to observe the structural specificity of the RNA granules. Third, while the sm-FRET analysis for longer RNAs remains challenging due to its distance limitations, investigation of the longer region of gRNA involving the multiple domains might be essential for the understanding of how the compaction of multiple granules are achieved. Thus, the current findings lay the foundation for new directions in the study of the genome packaging of SARS-CoV-2.

## ACKNOWLEDGMENTS

Y.I is grateful for JSPS KAKENHI Grants JP24K18075, and S.T. for JP20K21166, JP21H02438, and JP24K01983. S.T. is also grateful for JST CREST JPMJCR20H8. This work was performed under the Cooperative Research Program of “NJRC Mater. & Dev. (MEXT).” A.N.N., and S.T. acknowledge the support from the India Japan Science and Technology Cooperation Programme grant no. DST/INT/JSPS/P-394/2024 (G).

## References

1. E. Hartenian, et al., The molecular virology of coronaviruses. Journal of Biological Chemistry 295, 12910–12934 (2020).

2. S. Steiner, et al., SARS-CoV-2 biology and host interactions. Nat Rev Microbiol 22, 206–225 (2024).

3. P. V’kovski, A. Kratzel, S. Steiner, H. Stalder, V. Thiel, Coronavirus biology and replication: implications for SARS-CoV-2. Nat Rev Microbiol 19, 155–170 (2021).

4. E. Grellet, I. L’Hôte, A. Goulet, I. Imbert, Replication of the coronavirus genome: A paradox among positive-strand RNA viruses. Journal of Biological Chemistry 298, 101923 (2022).

5. D. Bracquemond, D. Muriaux, Betacoronavirus Assembly: Clues and Perspectives for Elucidating SARS-CoV-2 Particle Formation and Egress. mBio 12, 10.1128/mbio.02371-21 (2021).

6. W. Wu, Y. Cheng, H. Zhou, C. Sun, S. Zhang, The SARS-CoV-2 nucleocapsid protein: its role in the viral life cycle, structure and functions, and use as a potential target in the development of vaccines and diagnostics. Virol J 20, 6 (2023).

7. W. Yan, Y. Zheng, X. Zeng, B. He, W. Cheng, Structural biology of SARS-CoV-2: open the door for novel therapies. Sig Transduct Target Ther 7, 26 (2022).

8. H. Yang, Z. Rao, Structural biology of SARS-CoV-2 and implications for therapeutic development. Nat Rev Microbiol 19, 685–700 (2021).

9. P. S. Masters, Coronavirus genome packaging and nucleocapsid assembly. Journal of Virology 100, e01330–25 (2026).

10. J. D. Pata, T. K. Stevens, L. Kuo, P. S. Masters, Definition of the components required for selective packaging of coronavirus genomic RNA. Proc. Natl. Acad. Sci. U.S.A. 122, e2513552122 (2025).

11. M. Morse, J. Sefcikova, I. Rouzina, P. J. Beuning, M. C. Williams, Structural domains of SARS-CoV-2 nucleocapsid protein coordinate to compact long nucleic acid substrates. Nucleic Acids Research 51, 290–303 (2023).

12. S. Klein, et al., SARS-CoV-2 structure and replication characterized by in situ cryo-electron tomography. Nat Commun 11, 5885 (2020).

13. H. Yao, et al., Molecular Architecture of the SARS-CoV-2 Virus. Cell 183, 730–738.e13 (2020).

14. C. R. Carlson, et al., Reconstitution of the SARS-CoV-2 ribonucleosome provides insights into genomic RNA packaging and regulation by phosphorylation. Journal of Biological Chemistry 298, 102560 (2022).

15. N. Kaneda, et al., A Single Dimer of the SARS-CoV-2 N Protein Can Associate with Multiple Fragments of Single-Stranded and Stem-Loop RNA: A Single-Molecule FRET and FCS Investigation. ACS Omega 11, 21382–21394 (2026).

16. G. Wolff, S. Zheng, A. J. Koster, E. J. Snijder, M. Bárcena, A molecular pore spans the double membrane of the coronavirus replication organelle. Science (New York, N.Y.) 369, 1395–1398 (2020).

17. A. N. Adly, et al., Assembly of SARS-CoV-2 ribonucleosomes by truncated N∗ variant of the nucleocapsid protein. Journal of Biological Chemistry 299 (2023).

18. C. R. Carlson, et al., Phosphoregulation of Phase Separation by the SARS-CoV-2 N Protein Suggests a Biophysical Basis for its Dual Functions. Molecular Cell 80, 1092–1103.e4 (2020).

19. C. Cao, et al., The architecture of the SARS-CoV-2 RNA genome inside virion. Nat Commun 12, 3917 (2021).

20. H. Chen, et al., Liquid–liquid phase separation by SARS-CoV-2 nucleocapsid protein and RNA. Cell Res 30, 1143–1145 (2020).

21. N. C. Kathe, M. Novakovic, F. H.-T. Allain, J. Woodruff, Buffer choice and pH strongly influence phase separation of SARS-CoV-2 nucleocapsid with RNA. Molecular Biology of the Cell 35, ar73 (2024).

22. M. Dang, Y. Li, J. Song, ATP biphasically modulates LLPS of SARS-CoV-2 nucleocapsid protein and specifically binds its RNA-binding domain. Biochemical and Biophysical Research Communications 541, 50–55 (2021).

23. S. M. Cascarina, E. D. Ross, Phase separation by the SARS-CoV-2 nucleocapsid protein: Consensus and open questions. Journal of Biological Chemistry 298, 101677 (2022).

24. C. Iserman, et al., Genomic RNA Elements Drive Phase Separation of the SARS-CoV-2 Nucleocapsid. Molecular Cell 80, 1078–1091.e6 (2020).

25. S. Zúñiga, et al., Coronavirus nucleocapsid protein is an RNA chaperone. Virology 357, 215–227 (2007).

26. P. R. Bezerra, F. C. L. Almeida, Structural basis for the participation of the SARS-CoV-2 nucleocapsid protein in the template switch mechanism and genomic RNA reorganization. Journal of Biological Chemistry 300, 107834 (2024).

27. L. Zargarian, et al., Chaperone Activity of SARS-CoV-2 Nucleocapsid Protein: RNA Annealing and Destabilization Mechanisms. Journal of Molecular Biology 437, 169312 (2025).

28. B. Zhang, et al., The regulatory mechanisms of SARS-CoV-2 N protein helicase and its annealing activity. iScience 28, 113983 (2025).

29. S. Babl, et al., SARS-COV-2 nucleocapsid protein variants have differential RNA chaperone activity. The FEBS Journal febs.70329 (2025). 10.1111/febs.70329.

30. T. Ha, C. Kaiser, S. Myong, B. Wu, J. Xiao, Next generation single-molecule techniques: Imaging, labeling, and manipulation in vitro and in cellulo. Molecular Cell 82, 304–314 (2022).

31. A. Joshi, et al., Single-molecule FRET unmasks structural subpopulations and crucial molecular events during FUS low-complexity domain phase separation. Nat Commun 14, 7331 (2023).

32. D. Nettels, et al., Single-molecule FRET for probing nanoscale biomolecular dynamics. Nat Rev Phys 6, 587–605 (2024).

33. J. Sankaran, T. Wohland, Current capabilities and future perspectives of FCS: super-resolution microscopy, machine learning, and in vivo applications. Commun Biol 6, 699 (2023).

34. J. Enderlein, I. Gregor, D. Patra, T. Dertinger, U. B. Kaupp, Performance of Fluorescence Correlation Spectroscopy for Measuring Diffusion and Concentration. ChemPhysChem 6, 2324–2336 (2005).

35. M. Z. Palo, et al., Conserved long-range interactions are required for stable folding of orthoflaviviral genomic RNA. Nucleic Acids Res 53, gkaf514 (2025).

36. W.-J. C. Lai, et al., mRNAs and lncRNAs intrinsically form secondary structures with short end-to-end distances. Nat Commun 9, 4328 (2018).

37. Z. Xie, N. Srividya, T. R. Sosnick, T. Pan, N. F. Scherer, Single-molecule studies highlight conformational heterogeneity in the early folding steps of a large ribozyme. Proc. Natl. Acad. Sci. U.S.A. 101, 534–539 (2004).

38. N. C. Huston, et al., Comprehensive in vivo secondary structure of the SARS-CoV-2 genome reveals novel regulatory motifs and mechanisms. Molecular Cell 81, 584–598.e5 (2021).

39. K. Kuwahara, S. Yajima, Y. Yamano, F. Nagatsugi, K. Onizuka, Formation of Direction-Controllable Pseudorotaxane and Catenane Using Chemically Cyclized Oligodeoxynucleotides and Their Noncovalent RNA Labeling. Bioconjugate Chem. (2023).

40. G. J. Smith, T. R. Sosnick, N. F. Scherer, T. Pan, Efficient fluorescence labeling of a large RNA through oligonucleotide hybridization. RNA 11, 234–239 (2005).

41. C. Mirkes, Š. Zbýň, G. L. Chadzynski, S. Trattnig, K. Scheffler, “MRI using 23Na” in Encyclopedia of Spectroscopy and Spectrometry, (Elsevier, 2017), pp. 911–918.

42. N. K. Lee, et al., Accurate FRET Measurements within Single Diffusing Biomolecules Using Alternating-Laser Excitation. Biophysical Journal 88, 2939–2953 (2005).

43. A. N. Kapanidis, et al., Fluorescence-aided molecule sorting: Analysis of structure and interactions by alternating-laser excitation of single molecules. Proceedings of the National Academy of Sciences 101, 8936–8941 (2004).

44. S. Mitra, et al., Flexible Target Recognition of the Intrinsically Disordered DNA-Binding Domain of CytR Monitored by Single-Molecule Fluorescence Spectroscopy. J. Phys. Chem. B 126, 6136–6147 (2022).

45. Z.-J. Tan, S.-J. Chen, RNA Helix Stability in Mixed Na+/Mg2+ Solution. Biophysical Journal 92, 3615–3632 (2007).

46. A. S. Petrov, J. C. Bowman, S. C. Harvey, L. D. Williams, Bidentate RNA–magnesium clamps: On the origin of the special role of magnesium in RNA folding. RNA 17, 291–297 (2011).

47. Q. Hou, S. Chatterjee, P. E. Lund, K. C. Suddala, N. G. Walter, Single-molecule FRET observes opposing effects of urea and TMAO on structurally similar meso- and thermophilic riboswitch RNAs. Nucleic Acids Research 51, 11345–11357 (2023).

48. A. Plumridge, K. Andresen, L. Pollack, Visualizing Disordered Single-Stranded RNA: Connecting Sequence, Structure, and Electrostatics. J. Am. Chem. Soc. 142, 109–119 (2020).

49. A. Borodavka, et al., Sizes of Long RNA Molecules Are Determined by the Branching Patterns of Their Secondary Structures. Biophysical Journal 111, 2077–2085 (2016).

50. M. A. Digman, R. Dalal, A. F. Horwitz, E. Gratton, Mapping the Number of Molecules and Brightness in the Laser Scanning Microscope. Biophysical Journal 94, 2320–2332 (2008).

51. A. M. P. Romani, Cellular magnesium homeostasis. Archives of Biochemistry and Biophysics 512, 1–23 (2011).

52. A. Chen, et al., A coronaviral pore-replicase complex links RNA synthesis and export from double-membrane vesicles. Sci. Adv. 10, eadq9580 (2024).

53. E. Murigneux, et al., Proteomic analysis of SARS-CoV-2 particles unveils a key role of G3BP proteins in viral assembly. Nat Commun 15, 640 (2024).

54. B. Malone, N. Urakova, E. J. Snijder, E. A. Campbell, Structures and functions of coronavirus replication–transcription complexes and their relevance for SARS-CoV-2 drug design. Nat Rev Mol Cell Biol 23, 21–39 (2022).

55. Y. Zhang, et al., In vivo structure and dynamics of the SARS-CoV-2 RNA genome. Nat Commun 12, 5695 (2021).

56. L. Sun, et al., In vivo structural characterization of the SARS-CoV-2 RNA genome identifies host proteins vulnerable to repurposed drugs. Cell 184, 1865–1883.e20 (2021).

57. J. Vieregg, W. Cheng, C. Bustamante, I. Tinoco, Measurement of the Effect of Monovalent Cations on RNA Hairpin Stability. J. Am. Chem. Soc. 129, 14966–14973 (2007).

58. Ignacio Tinoco, Carlsom Bustamante, How RNA folds. Journal of Molecular Biology 293, 271–81 (1999).

59. D. E. Draper, A guide to ions and RNA structure. RNA 10, 335–343 (2004).

60. A. Martinez-Monge, I. Pastor, C. Bustamante, M. Manosas, F. Ritort, Measurement of the specific and non-specific binding energies of Mg2+ to RNA. Biophysical Journal 121, 3010–3022 (2022).

61. A. M. Yoffe, et al., Predicting the sizes of large RNA molecules. Proc. Natl. Acad. Sci. U.S.A. 105, 16153–16158 (2008).

62. A. Rastandeh, et al., Measuring the selective packaging of RNA molecules by viral coat proteins in cells. Proc. Natl. Acad. Sci. U.S.A. 122, e2505190122 (2025).

63. J. A. Imperatore, et al., Highly conserved s2m element of SARS-CoV-2 dimerizes via a kissing complex and interacts with host miRNA-1307-3p. Nucleic Acids Research 50, 1017–1032 (2022).

64. K. K. Sharma, et al., Structural flexibility in the ordered domain of the dengue virus strain 2 capsid protein is critical for chaperoning viral RNA replication. Cell. Mol. Life Sci. 82, 184 (2025).

65. F. Baudin, C. Bach, S. Cusack, R. W. Ruigrok, Structure of influenza virus RNP. I. Influenza virus nucleoprotein melts secondary structure in panhandle RNA and exposes the bases to the solvent. EMBO J 13, 3158–3165 (1994).

66. J. P. K. Bravo, et al., Structural basis of rotavirus RNA chaperone displacement and RNA annealing. Proc. Natl. Acad. Sci. U.S.A. 118, e2100198118 (2021).

67. X. E. Yong, V. R. Palur, G. S. Anand, T. Wohland, K. K. Sharma, Dengue virus 2 capsid protein chaperones the strand displacement of 5′-3′ cyclization sequences. Nucleic Acids Res 49, 5832–5844 (2021).

68. D. Herschlag, RNA Chaperones and the RNA Folding Problem. Journal of Biological Chemistry 270, 20871–20874 (1995).

69. J. G. Levin, J. Guo, I. Rouzina,, K. Musier-Forsyth, “Nucleic Acid Chaperone Activity of HIV-1 Nucleocapsid Protein: Critical Role in Reverse Transcription and Molecular Mechanism” in Progress in Nucleic Acid Research and Molecular Biology, (2005), pp. 217–286.

70. P. M. Laughlin, K. Young, G. Gonzalez-Gutierrez, J. C.-Y. Wang, A. Zlotnick, A narrow ratio of nucleic acid to SARS-CoV-2 N-protein enables phase separation. Journal of Biological Chemistry 300, 107831 (2024).

71. S. Lu, et al., The SARS-CoV-2 nucleocapsid phosphoprotein forms mutually exclusive condensates with RNA and the membrane-associated M protein. Nat Commun 12, 502 (2021).

72. C. A. Roden, et al., Double-stranded RNA drives SARS-CoV-2 nucleocapsid protein to undergo phase separation at specific temperatures. Nucleic Acids Research 50, 8168–8192 (2022).

73. B. Favetta, et al., Phosphorylation toggles the SARS-CoV-2 nucleocapsid protein between two membrane-associated condensate states. Nat Commun 16, 7970 (2025).

74. T. M. Perdikari, et al., SARS-CoV-2 nucleocapsid protein phase-separates with RNA and with human hnRNPs. The EMBO Journal 39, e106478 (2020).

