## Supplementary Information for "A Structural Domain in the genomic RNA of SARS-CoV-2 Folds into a Compact Granular Structure without the N protein: A Single-Molecule Fluorescence Spectroscopic Investigation"

### Supporting Text

#### Preparation of $gRNA_{12k}$ corresponding to nt 12230-12686 of SARS-CoV-2 genomic RNA (gRNA)

A template DNA sequence for tag-$gRNA_{12k}$ was inserted into a plasmid, pEX-A2J2 by Eurofins Genomics (Supporting Table S1). The DNA contains the T7 promotor sequence, an additional tag sequence and the sequence from nt 12230 to 12686 of gRNA. The 5′-tag sequence was taken from a previous study (1). The DNA fragment was amplified by PCR with Prime STAR Max premix (Takara Bio) using the forward and reverse primers (Supporting Table S1). The PCR product was loaded onto 2% Tris-acetate-EDTA agarose gel and run at 50 V for 1.5 h. The PCR products stained with SYBR gold (Thermo Fisher) were excised from the gel, followed by the purification with Monarch DNA Gel Extraction Kit (New England Biolabs). To prepare tag-$gRNA_{12k}$, we performed *in vitro* transcription using an RNA Synthesis Kit (HiScribe T7 Quick High Yield, New England Biolabs). Briefly, the purified PCR product was mixed with T7 RNA Polymerase Mix and NTP Buffer Mix (New England Biolabs), and was incubated at 37°C for ~16 hours. The DNaseI was added to the mixture, followed by the purification with Monarch Spin RNA Cleanup Kit (New England Biolabs). The absorbance at 260 nm was measured to estimate the concentration of $gRNA_{12k}$.

#### Labeling of fluorophores

We labeled pCp-AZdye488 or pCp-AZdye647 into the 3′ end of tag-$gRNA_{12k}$ using ligase I. The tag-$gRNA_{12k}$ was mixed at a concentration of 10–14 μM with 1x reaction buffer (New England Biolab), 1 mM ATP (New England Biolab) and 10% DMSO (FUJIFILM Wako Pure Chemical). The mixture was incubated at 65°C for 5 min, followed by an incubation at room temperature for 3 min. In some cases, 50% polyethylene glycol 8000 (PEG8000, New England Biolab) was added to the mixture. To the mixture, 1.67 unit/μL ligase I (New England Biolabs) and 50–66 μM pCp-AZdye488 or pCp-AZdye647 was added followed by an incubation at 16°C for 16 hours. Finally, tag-$gRNA_{12k}$ reacted with pCp-AZdye488 (tag-$gRNA_{12k}$–488) or pCp-AZdye647 (tag-$gRNA_{12k}$–647) were purified with Monarch RNA clean up kit (New England Biolabs). If PEG8000 was added in the reaction mixture, the labeled sample was purified with the same kit twice.

We purchased 22 nt 2′-*O*-methylated oligonucleotide with a sequence complementary to the 5′-tag sequence labeled with Alexa647 or Alexa488 at the 5′ end (2′-OMe-647 or 2′-OMe-488). The 2′-OMe-647 or 2′-OMe-488 were hybridized into the 5′ tag region of tag-gRNA$_{12k}$–488 or tag-gRNA$_{12k}$–647, respectively (2). tag-gRNA$_{12k}$–488 was incubated at 90°C for 2 min in 20 mM Tris-HCl buffer adjusted at pH 8.1, followed by an incubation for 3 min at room temperature. A solution containing NaCl was added to the final concentration of 50 mM, and the resultant mixture was heated to 37°C for 5 min. 2′-OMe-647 was then added and the sample was incubated at 37°C for 1 hour. The final concentrations of tag-gRNA$_{12k}$–488 and 2′-OMe-647 in the hybridizing solution were equal, ranging from 1.3 to 2.0 μM. The resultant sample was named as 647=tag-gRNA$_{12k}$–488. Similarly, 2′-OMe-488 was hybridized into the 5′ tag of tag-gRNA$_{12k}$–647 and named as 488=tag-gRNA$_{12k}$–647. To prepare the sample singly labeled with Alexa488, we hybridized non-labeled 2′-OMe oligonucleotide with the 5′-tag of tag-gRNA$_{12k}$–488 and named as 2′-OMe=tag-gRNA$_{12k}$–488.

After the hybridization, the labeled samples were purified by using agarose gel electrophoresis and a purification kit. The labeled samples were applied into 2% agarose gel and subjected to the electrophoresis. The gel was then imaged using an excitation light at 480–520 nm with illuminator (NIPPON Genetics Co, Ltd), and a region of the gel corresponding to the labeled samples was extracted and purified with Monarch spin RNA clean up kit. The concentration of the double labeled samples was estimated by converting their absorbance at 260 nm assuming that $A_{260}$ = 1.0 corresponds to 40 μg/mL. The labeling efficiencies of Alexa488 to the 3′ end of the tag-gRNA$_{12k}$ and Alexa647 to the 5′ end were roughly estimated to be ~80% and ~60%, respectively, using the absorbance at 260 nm, 493 nm and 651 nm.

**Expression and purification of the SARS-CoV-2 N protein**

All chemicals used for the sample preparation and the spectroscopic measurements were commercially available. The SARS-CoV-2 N protein from the original Wuhan variant, whose N-terminus is tagged with 10 histidines, was constructed as previously reported (3). A single glycine residue was added to the N-terminus of the His-tag. The N protein was expressed and purified as previously described (3). Briefly, the gene for the N protein was cloned into the pET-28a(+) vector by Eurofins Genomics, which was transformed into *E. coli*, BL21(DE3). The cells were grown at 37°C until the optical density at 600 nm reached ~0.6, and were further cultured for 4 h after adding 1 mM isopropyl β-D-1-thiogalactopyranoside. The cells were harvested and disrupted by sonication, and the solution was centrifuged. The pellet was washed three times with the buffer containing 20 mM Tris-HCl at pH 8.0, 200 mM NaCl, 10–20 μg/mL ribonuclease A and 0.3% Tween-20. The pellet was resuspended overnight with the buffer containing 20 mM Tris-HCl at pH 7.5, 1 M NaCl and 6 M urea. The N protein was purified by using a Ni-NTA column (EconoFit Nuvia IMAC Column 1 mL, Bio-Rad) installed in the NGC Chromatography System (Bio-Rad). The N protein was eluted and passed through a desalting column (PD MiniTrap G-25, Cytiva) equilibrated with the storage buffer containing 20 mM Tris-HCl at pH 7.5 and 1 M NaCl . The concentration of the N protein was calculated by measuring the absorbance at 280 nm using $\varepsilon = 43890\ \mathrm{L\cdot mol^{-1}\cdot cm^{-1}}$.

**Sample preparation for the FCS or sm-FRET measurements**

The protocol for the preparation of the samples for the FCS and sm-FRET measurements was as follows. The labeled RNAs were incubated at 20 nM in 20 mM Tris-HCl at pH 7.5 for 5 min at either 37°C or 75°C, followed by an incubation at 37°C for 5 min. As necessary, the non-labeled tag-$gRNA_{12k}$ was added to the labeled samples before the incubation. Next, either an arbitrary concentration of cations or the N protein or both was added to the labeled RNA solution, resulting in the final concentration of the labeled RNA from 50 to 100 pM or from 1 to 2 nM for sm-FRET and FCS measurements, respectively.

The final mixture was incubated at 37°C for 10 min and transferred to glass base dishes for the spectroscopic measurements at room temperature (~22°C).

**Preparation of 647=tag-gRNA$_{12k}$–488 with ASOs for the sm-FRET measurements**

Solutions containing 20 mM Tris-HCl buffer at pH 8.1, 20-nM 647=tag-gRNA$_{12k}$–488 and 0-, 2.0-, 20- or 200-nM ASO were prepared, heated at 75°C for 5 min to unfold the sample without releasing 2′-OMe-647 and incubated at 37°C for 5 min. Subsequently, an equal volume of 150 mM NaCl in 20 mM Tris-HCl buffer was added to the final concentrations of 647=tag-gRNA$_{12k}$–488 at 10 nM, ASO at 0, 1.0, 10 or 100 nM and NaCl at 75 mM, and incubated at 37°C for one hour. After the incubation, the solutions were diluted for 125 times using 20 mM Tris-HCl buffer at pH 8.1 containing 150 mM NaCl, and used for the sm-FRET measurements. A precise control of the NaCl concentration during the incubation period was important to replace the long-range base pairing with ASOs. In the absence of NaCl during the incubation, the replacement of the long-range base pairing with ASO2 did not occur likely due to the intermolecular electrostatic repulsion between 647=tag-gRNA$_{12k}$–488 and ASO2 (Supporting Fig. S3). The addition of 75 mM NaCl during the incubation led to the hybridization of ASOs before the formation of the long-range base pairing in a manner dependent on the ASO concentration (Fig. 3 and Supporting Fig. S3). However, the disruption of the long-range base pairing did not occur in the presence of 100 nM of rA$_{13}$ during the incubation. The addition of 750 mM NaCl during the incubation likely stabilized the intramolecular long-range base pair (Fig. 2B), and accordingly, the formation of the intermolecular base pair between 647=tag-gRNA$_{12k}$–488 and ASO2 became less likely, resulting in the remaining of the high-efficiency peak even in the presence of ASO2 (Supporting Fig. S3).

**Detection system for the sm-FRET measurements and for the fluorescence intensity fluctuation measurements at the 642-nm excitation**

We performed the sm-FRET measurements by using a home-made confocal optics equipped with the Alternating Laser Excitation (ALEX) system reported previously (3). In brief, the 488 nm laser for the excitation of donor (OBIS 488 nm LX 100 mW, 1236444, Coherent) and 642 nm laser for the excitation of acceptor (OBIS 640 nm LX 75 mW, 1236445, Coherent) was introduced to a water immersion objective (NA 1.2, CFI PlanApo 60XC WI, Nikon), and focused into the sample placed on the glass-based dish (IWAKI). Fluorescence photons were collected by the same objective, passed through a dual bandpass filter (ZET488/640m, Chroma Technology), and were separated into donor and acceptor photons by a dichroic mirror (593 nm cut-on wavelength, #67-083, Edmund Optics). The separated photons were passed through bandpass filters (for donor, FBH520-40, Thorlabs, and for acceptor, FF01-676/29, Semrock), and detected by two single-photon avalanche diodes (SPADs, SPCM AQRH-14-FC, Excelitas). Signals from SPADs were recorded as photon arrival times by using a counter (PCIe-6612, National Instruments) operated by a homebuilt LabVIEW software (National Instruments) and stored in the photon-HDF5 file format (4). The laser power was measured near the focus point of the objective, and were adjusted to ~40 μW and ~20 μW for the donor and acceptor excitations, respectively. The same optical system was used for the fluorescence intensity fluctuation measurements at 642-nm excitation. We used the 642 nm laser and adjusted its power at the sample point to ~10 μW. The time interval for the photon intensity measurements was set at 10 ms.

For the ALEX and fluorescence fluctuation measurements, the inner surface of glass base dishes (3971-035, AGC Techno Glass) was coated with 2-methacryroyloxyethylphosphorylcholine (MPC) polymer (Lipidure CM5206, NOF) by rinsing the surface by 0.5% (w/v) solution of MPC in 99.5% ethanol (052-06925, Fujifilm Wako Pure Chemical) followed by drying. 70–180 μL of the sample solution was placed on the coated dish. Before measuring the RNA data, we measured 20–40 pM of 38 bp dsDNA in which Alexa488 and Alexa647 were labeled with the 23rd thymine base from the 5′ end in one strand and with the 31st thymine base from the 5′ end in the complementary strand, respectively (5), to calibrate the ALEX apparatus. The concentrations of the labeled RNA samples for the ALEX measurements were between 50 pM and 100 pM which were determined based on the RNA concentration. The concentration of the labeled RNA samples for the fluorescence fluctuation measurements was 2 nM. In the ALEX and fluorescence fluctuation measurements, the data acquisition period was typically 60 min and 10 min, respectively. All measurements were conducted in a room whose temperature was adjusted at ~22°C.

**ALEX data analysis**

The analysis of the ALEX data was based on the standard protocol using the FRETBursts software as described previously (6). First, the local background rates of the four photon types, the donor photon during donor excitation, the acceptor photon during donor excitation, the donor photon during acceptor excitation and the acceptor photon during acceptor excitation, were calculated in windows of 30 s. Second, the fluorescence bursts were searched for each photon types by selecting the time regions where the count rate calculated using arrival times of 10 consecutive photons exceeded the count rate threshold set to 6 times the local background rates. Third, bursts were selected from the searched bursts by using the burst size threshold of 20 for the sum of donor and acceptor photons obtained during the donor excitation to eliminate the acceptor-only species. Fourth, another burst selection was performed by setting the burst size threshold of 20 for the acceptor photons collected during the acceptor excitation to eliminate the donor-only bursts. From the photon counts in each channel, apparent FRET efficiency (*E*) and stoichiometry (*S*) for each burst were calculated based on eq. S1 and S2:

$$E = \frac{F_{Dex}^{Aem}}{\gamma F_{Dex}^{Dem} + F_{Dex}^{Aem}} \qquad \text{eq. S1}$$

$$S = \frac{\gamma F_{Dex}^{Dem} + F_{Dex}^{Aem}}{\gamma F_{Dex}^{Dem} + F_{Dex}^{Aem} + F_{Aex}^{Aem}} \qquad \text{eq. S2}$$

where $F_{Dex}^{Aem}$ is the number of acceptor emission photons during the donor excitation in one burst. The subscripts *Dex* and *Aex* denote the donor and acceptor excitation, respectively, and the superscript *Dem* and *Aem* denote the donor and acceptor emission, respectively. The $\gamma$ value represents the ratio of the florescence detection efficiencies for the donor and acceptor channels and was set to the estimated value of 0.48 (7). *S* depends on the intensities of the two lasers used to excite the donor and acceptor.

**Detection system for the FCS measurements based on the 484-nm excitation**

The FCS measurements based on the 484-nm excitation was conducted using a handmade FCS spectrometer reported previously (3). In brief, 484-nm laser light (LDS1003, Precise Gauges) was focused onto the sample solution using a water immersion objective (UPlanSApo 60x, Olympus). The fluorescence photons were collected by the same objective, passed through two dichroic mirrors (P50H, Thorlabs, and 67-083, Edmund), a bandpass filter (FBH520-40, Thorlabs) and detected by HPD (H13223-40, Hamamatsu Photonics). The HPD outputs were recorded using a time correlated single photon counting module (TimeHarp 260 NANO, PicoQuant). The photon data were converted into the photon intensity correlations using SymPhoTime64 software (PicoQuant). The FCS measurements based on the 642 nm excitation was conducted using the same optics developed for the ALEX measurements. A single unit of SPAD was used to detect all the fluorescence photons of Alexa647. The fluorescence correlations were calculated by using a home-made software.

The glass base dishes coated with the MPC polymer were used for the FCS measurements. 70–80 μL of the sample solution was placed on the dish and covered with a lid. The FCS measurements were performed in a room adjusted at 22°C. The excitation laser power at 488 nm was measured near the focus point of the objective, and was adjusted to 50 μW. The data acquisition time for one measurement was 10 min. Solutions containing 1.0–2.0 nM of the labeled samples were prepared and measured. The sample concentrations were based on the RNA concentration. Solutions containing 1 nM rhodamine 110 in water were measured once per day as a reference. Moreover, a solution containing 0.5–1.0 nM pCp-AZdye488

(NU-1706-488, Jena BioScience) was also measured once per day whose diffusion times were used for the FCS curve fitting.

**Fitting of the correlation function and estimation of $R_H$**

To estimate $R_H$ of OMe=tag-gRNA$_{12k}$–488, we first fitted the correlograms of the reference samples in the time domain from 1.0 × $10^{-6}$ s to 1.0 s with eq. S3 using fitting algorithm of Igor Pro 9:

$$G(\tau) = G(0) \cdot \frac{1 - frac + frac \cdot \exp(-\frac{\tau}{\tau_F})}{1 - frac} \cdot \frac{1}{\left(1 + \frac{\tau}{\tau_D}\right) \cdot \left(1 + \frac{\tau}{\tau_D \cdot s^2}\right)^{-\frac{1}{2}}} + G(\infty) \qquad \text{eq. S3}$$

where $frac$, $\tau_F$, $\tau_D$ and $s$ were the ratio of triplet state contribution, the triplet lifetime, the diffusion time of the fluorophores, the ratio of the axial radius to the radial radius of the observation volume, respectively. $G(\infty)$ was set to 0. The $s$ value was calculated from the correlation function of rhodamine 110 and used as the fixed parameter in the fitting of other samples obtained in the same day.

Following our previous study (3), the correlation functions of the labeled samples, $G(\tau)$, were fit with eq. S4, assuming the presence of two diffusing components having different diffusivity and brightness: one is the free fluorophore diffusing quickly with a constant brightness and the other is the RNA samples labeled with Alexa488 diffusing slowly with a variable brightness:

$$G(\tau) = \frac{b^2}{\langle I^2 \rangle} \cdot \frac{1 - frac + frac \cdot \exp(-\frac{\tau}{\tau_F})}{1 - frac}$$

$$\cdot \left\{ \left( \frac{< I > -bkg}{b} - a \cdot n_R \right) \cdot \frac{1}{\left(1 + \frac{\tau}{\tau_A}\right) \cdot \left(1 + \frac{\tau}{\tau_A \cdot s^2}\right)^{-\frac{1}{2}}} + a^2 \cdot n_R \right.$$

$$\left. \cdot \frac{1}{\left(1 + \frac{\tau}{\tau_R}\right) \cdot (1 + \frac{\tau}{\tau_R \cdot s^2})^{-\frac{1}{2}}} \right\} + G(\infty) \qquad \text{eq. S4}$$

where $b$, $<I>$, $bkg$, $a$, $n_R$, $\tau_A$, $\tau_R$ are the brightness of free Alexa488 or Alexa647, the averaged fluorescence intensity during the FCS measurement, the background photon count, the ratio of the brightness of the labeled RNA relative to that of the free fluorophore and the number of the labeled RNA in the observation volume, the translational correlation time for the free fluorophore, and the translational correlation time of the labelled RNA, respectively. $<I>$ can be expressed as follows:

$$\langle I \rangle = b \cdot (n_A + a \cdot n_R) + bkg \qquad \text{eq. S5}$$

where $n_A$ is the number of the free fluorophore in the observation volume. The brightness of free Alexa488 and Alexa647 is assumed to be the same as that of RNA-labeled dyes in the absence of the N protein, that is, $a = 1$ (Supporting Tables S2 and S4). We fitted the correlograms in the absence of the N protein using eq. S4, to estimate $n_R$ and $\tau_R$. In the presence of the N protein, we fixed the $b$ obtained in the absence of the N protein to estimate $a$, $n_R$ and $\tau_R$. The reduction in $a$ might be due to the quenching of the Alexa488 by the N protein.

We calculated the hydrodynamic radius ($R_H$) of the labelled RNA by converting $\tau_R$ estimated by the fitting to a diffusion coefficient ($D_R$) using eq. S6, and converting $D_R$ to $R_H$ using eq. S7:

$$D_R = \frac{{\omega_0}^2}{4 \cdot \tau_R} \qquad \text{eq.}\, S6$$

$$R_{\mathrm{H}} = \frac{k_B \cdot T}{6 \cdot \pi \cdot \eta \cdot D_R} \qquad \text{eq.}\, S7$$

where $k_{\mathrm{B}}$ is the Boltzmann constant ($1.38 \cdot 10^{-23}$ J/K), $T$ is the absolute temperature (293 K) and $\eta$ is the viscosity for water at 293 K in the presence of 150 mM NaCl ($1.016 \cdot 10^{-3}$ Pa·s) or in the absence of NaCl ($1.002 \cdot 10^{-3}$ Pa·s) (8). The $\omega_0$ value is the short radius of the focus area, which was determined using the reference FCS data for the free rhodamine 110 whose diffusion coefficients was 440 $\mu m^2$/s (9).

As discussed above, determining $R_{\mathrm{H}}$ for long RNA samples even in the absence of the N protein involves multiple steps that may introduce experimental variance. These factors include the inherent heterogeneity of samples prepared via *in vitro* transcription, the necessity for a two-component fitting model to account for residual free dye, and the requirement for standard references measured under identical conditions. To ensure data reliability, we performed two independent FCS measurements for 2′-OMe=tag-gRNA$_{12k}$–488 and conducted separate fitting analyses. The final $R_{\mathrm{H}}$ values were determined by averaging the results of these two measurements. The detailed fitting parameters obtained in one measurement, and the averaged $R_{\mathrm{H}}$ based on the two measurements are provided in Supporting Tables S2 and S3, respectively. The averaged results were plotted in Supporting Figure S4.

The determination of $R_{\mathrm{H}}$ for 2′-OMe=tag-gRNA$_{12k}$–488 in the presence of N protein based on the FCS measurements excited at 484 nm proved more challenging due to several factors. First, the long RNA samples started to aggregate when the N protein concentration reached 100 nM. Second, the formation of a heterogeneous mixture containing multiple oligomeric states rendered the two-component fitting model inadequate. Third, sample conditions fluctuated over time following the N protein addition, likely due to the adsorption of aggregates onto the cell surfaces. Fourth, the fluorescence of Alexa488 labeled to RNA was significantly quenched upon the N protein binding, leading to substantial signal reduction. The $R_H$ values were determined by averaging the results of three independent measurements. The detailed fitting parameters obtained in one measurement, and the averaged $R_H$ based on the three measurements are provided in Supporting Tables S4 and S5, respectively. The averaged results were plotted in Figure 5B in the main text. To circumvent these issues, we changed our sample to 488=tag-gRNA$_{12k}$–647, in which Alexa647 was conjugated to the 3' end. After 488=tag-gRNA$_{12k}$–647 were mixed with N protein or buffer for at least 25 min, fluorescence intensity fluctuation measurements at 642 nm excitation were performed and the data were analyzed using the brightness and number method as described below.

**Brightness and number analysis of the fluorescence fluctuation data**

At the 642-nm excitation, we monitored the fluorescence fluctuations of 488=tag-gRNA$_{12k}$–647 at a sampling interval of 10 ms, and conducted the "brightness and number" (B&N) analysis (10–12). We define the fluorescence intensity time series, $I(t)$, as the sum of a constant background ($bkg$) and the emission from RNA particles ($I_{RNA}(t)$), whose number ($n_R$) fluctuates within the 10 ms time interval. The constant background includes both instrumental noise and emission from free fluorophores in the sample, which diffuse rapidly enough to contribute only as a baseline offset at this time scale. The brightness of the particle is denoted as $\varepsilon$ (equivalent to $b{\cdot}a$ in eq. S5). Since the variance of $I(t)$, $\sigma^2$, is primarily attributed to $I_{RNA}(t)$, $\varepsilon$ and $n_R$ can be derived from the following relationships:

$$\epsilon = \frac{\sigma^2}{\langle I_{sig}(t)\rangle} - 1 \qquad \text{eq. S8}$$

$$n_R = \frac{\langle I_{sig}(t)\rangle}{\epsilon} \qquad \text{eq. S9}$$

where $\langle I_{sig}(t)\rangle = \langle I(t)\rangle$ - *bkg* represents the mean background-subtracted intensity. This analysis remains valid for the heterogeneous emitting species. In such cases, the estimated parameters $\epsilon_{obs}$ and $n_{obs}$ represent weighted averages defined as follows:

$$\epsilon_{obs} = \frac{\sum n_i \cdot \epsilon_i{}^2}{\sum n_i \cdot \epsilon_i} \qquad \text{eq. S10}$$

$$n_{obs} = \frac{(\sum n_i \cdot \epsilon_i)^2}{\sum n_i \cdot \epsilon_i{}^2} \qquad \text{eq. S11}$$

For N protein concentrations of 0 nM and 10 nM, separate FCS measurements (data not shown) indicate that free fluorophores accounted for ~35% of the total fluorescence. For 100 nM and 1000 nM concentrations, where correlograms were not analyzable based on a two-component model, the background contribution was estimated at ~30%. This value was chosen to optimally reproduce the peak frequency of the observed photon counting histograms.

To validate the physical relevance of the estimated $\epsilon$ and $n_R$, we performed a model calculation to reconstruct the fluorescence intensity histograms. First, the input parameters $\epsilon$ and $n_R$ were determined from the background subtracted mean intensity, $\langle I_{sig}(t)\rangle = \langle I(t)\rangle$ - *bkg*, to isolate the contribution of the RNA particles. The model then integrates molecular number fluctuations, photon shot noise, and the spatial profile of the excitation volume. The occupancy of molecules $m$ within the effective observation volume $V_{eff}$ follows a Poisson distribution $P(m; n_R)$. Simultaneously, the photon detection from $m$ molecules follows a distribution $P_{sig}(k)$, which is the spatial average of the local Poissonian detection probabilities over the point spread function (PSF):

$$P_{sig}(k) = \frac{1}{V_{eff}} \int_V \mathrm{Pois}\left(k; m \cdot \epsilon_{peak} W(\mathbf{r})\right) d\mathbf{r} \qquad \text{eq. S12}$$

where $W(\mathbf{r})$ represents the normalized PSF profile and $\epsilon_{peak}$ is the maximum molecular brightness at the center of the focus. The relationship between the peak brightness $\epsilon_{peak}$ and the experimentally determined average brightness $\epsilon$ is governed by the shape factor $\gamma$:

$$\epsilon = \gamma \cdot \epsilon_{peak} \qquad \text{eq. S13}$$

We tried several values of $\gamma$ and adopted 0.7 that roughly reproduced the experimental histogram. To account for the experimental background noise, we modeled the background distribution $P_{\mathrm{bkg}}(k)$ as a Poisson distribution with a mean equal to the *bkg* value used in the B&N analysis. Finally, the theoretical histogram, $P_{total}(k)$, was generated by the convolution of $P_{sig}(k)$ and the background distribution $P_{bkg}(k)$:

$$P_{total}(k) = \left(P_{sig} * P_{bkg}\right)(k) = \sum_{j=0}^{k} P_{sig}(j) P_{bkg}(k-j) \qquad \text{eq. S14}$$

The semi quantitative agreement between the experimental histograms and the calculated models confirmed that our parameter estimation captured the behavior of the RNA molecules. It should be noted that this single-species model is specifically intended for systems with a uniform emitting population. In the case of heterogeneous samples, such as those observed in the presence of N protein at concentrations exceeding 100 nM, the experimental histograms became significantly broader, reflecting the presence of high-order aggregates with increased molecular brightness.

**CD measurements**

CD spectra of 0.5–0.8 μM $gRNA_{12k}$ in the buffer containing 20 mM Tris-HCl at pH 7.5 in the presence of different concentrations of NaCl, $MgCl_2$ and N protein were measured with a spectropolarimeter (J-720, Jasco) using a cell having 1 mm pathlength. The temperature dependency was measured using 1.0 μM $gRNA_{12k}$ in the buffer containing 20 mM Tris-HCl at pH 7.5 in the presence of 150 mM NaCl with a spectropolarimeter (J-815, Jasco) using a cell having 10 mm pathlength. Scanning speed was 10 nm/min.

### Supporting Tables

#### Supporting Table S1: The DNA and RNA sequences of the samples used in this investigation

| | |
|---|---|
| **The DNA for the preparation of tag-gRNA$_{12k}$[1)]** | 5' TAATACGACTCACTATAGGGAAAGCGGGCAGTGAGCGCAACG CAATTATCTGAATTTGACCGTGATGCAGCCATGCAACGTAAGTTG GAAAAGATGGCTGATCAAGCTATGACCCAAATGTATAAACAGGC TAGATCTGAGGACAAGAGGGCAAAAGTTACTAGTGCTATGCAGA CAATGCTTTTCACTATGCTTAGAAAGTTGGATAATGATGCACTCA ACAACATTATCAACAATGCAAGAGATGGTTGTGTTCCCTTGAAC ATAATACCTCTTACAACAGCAGCCAAACTAATGGTTGTCATACC AGACTATAACACATATAAAAATACGTGTGATGGTACAACATTTA CTTATGCATCAGCATTGTGGGAAATCCAACAGGTTGTAGATGCA GATAGTAAAATTGTTCAACTTAGTGAAATTAGTATGGACAATTC ACCTAATTTAGCATGGCCTCTTATTGTAACAGCTTTAAGGGCCAA TTCTGCTGTCAAATTACAGAAAATTA 3' |
| **Forward primer[2)]** | 5' TAATACGACTCACTATAGGGAAAGCG 3' |
| **Reverse primer[2)]** | 5' TAATTTTCTGTAATTTGACAGCAGAATTG 3' |
| **tag-gRNA$_{12k}$[3)]** | 5' gggaaagcgggcagugagcgcaacgcaauuaucugaauuugaccgugaugcagccaugcaac guaaguuggaaaagauggcugaucaagcuaugacccaaauguauaaacaggcuagaucugaggac aagagggcaaaaguuacuagugcuaugcagacaaugcuuuucacuaugcuuagaaaguuggaua augaugcacucaacaacauuaucaacaaugcaagagaugguuguguucccuugaacauaauaccu cuuacaacagcagccaaacuaaugguugucauaccagacuauaacacauauaaaaauacguguga ugguacaacauuuacuuaugcaucagcauugugggaaauccaacagguuguagaugcagauagu aaaauuguucaacuuagugaaauuaguauggacaauucaccuaauuuagcauggccucuuauug uaacagcuuuaagggccaauucugcugucaaauuacagaaaauua 3' |
| **2′-*O*-methyl-ribonucleotide[4)]** | 5' uugcguugcgcucacugcccgc 3' |
| **ASO1[5)]** | 5' ucuguaauuugac 3' |
| **ASO2[5)]** | 5' gucaaauucaga 3' |

[1)]The DNA sequence was amplified by PCR and inserted into a plasmid, pEX-A2J2. [2)]The forward and reverse primer sequences used for the PCR of the DNA sequences for constructing tag-gRNA$_{12k}$. The green letters correspond to a promoter sequence that was not transcribed. [3)]The blue letters correspond to regions proposed to form the long-range base pairs. The red letters correspond to the 5′ tag sequence for the labeling of Alexa647 by annealing the anti-sense sequence synthesized by using 2′-*O*-methylribonucleotides and Alexa647. Purple letters are additional sequences. The total number of nucleotides generated by the *in vitro* transcription was 494. [4)]The RNA sequences used for the labeling of Alexa647 or Alexa488 to the 5′ tag of tag-gRNA$_{12k}$. 2′-hydroxyl group of each nucleotide was methylated and either of the fluorophores was attached to the 5′ end of the 22-base sequence. The sequence is complementary to the 5′ tag region of tag-gRNA$_{12k}$. [5)]The RNA samples used in the antisense experiments. The sequences of ASO1 and ASO2 are complementary to the 3′ and 5′ regions of tag-gRNA$_{12k}$ assumed to form the long-range base pairing, respectively.

**Supporting Table S2: The parameters used in and obtained by the fitting analysis of a representative FCS data for 2′-OMe=tag-gRNA$_{12k}$-488 (1.5 nM) at various concentration of NaCl and $MgCl_2$**

| [NaCl] (mM)<br>[$MgCl_2$] (mM) | 0<br>0 | 0<br>0.1 | 0<br>1 | 0<br>10 | 150<br>0 | 150<br>0.1 | 150<br>1 | 150<br>10 |
|---|---|---|---|---|---|---|---|---|
| $s$[1)] | 7.5176 | 7.5176 | 7.5176 | 7.5176 | 7.5176 | 7.5176 | 7.5176 | 7.5176 |
| *frac*[2)] | 0.181 ± 0.006 | 0.181 ± 0.005 | 0.164 ± 0.008 | 0.160 ± 0.008 | 0.169 ± 0.008 | 0.205 ± 0.008 | 0.175 ± 0.008 | 0.184 ± 0.007 |
| $\tau_F$[3)] (μs) | 3.5 ± 0.2 | 3.8 ± 0.2 | 3.4 ± 0.2 | 4.0 ± 0.3 | 3.6 ± 0.3 | 3.4 ± 0.2 | 4.1 ± 0.3 | 4.4 ± 0.3 |
| $\tau_A$[4)] (μs) | 102.4 | 102.4 | 102.4 | 102.4 | 102.4 | 102.4 | 102.4 | 102.4 |
| $\tau_R$[5)] (μs) | 1063.1 ± 2.8 | 917.6 ± 2.5 | 1001.9 ± 2.9 | 951.8 ± 3.6 | 968.3 ± 3.5 | 868.6 ± 2.8 | 894.9 ± 3.6 | 847.3 ± 3.2 |
| $B$[6)] (photons/ms) | 16.88 ± 0.02 | 18.08 ± 0.02 | 12.95 ± 0.02 | 12.93 ± 0.02 | 12.58 ± 0.02 | 13.02 ± 0.02 | 13.11 ± 0.02 | 13.10 ± 0.02 |
| $I$[7)] (photons/ms) | 7.2 | 6.7 | 5.7 | 4.3 | 4.5 | 4.7 | 3.8 | 3.9 |
| $BG$[8)] (photons/ms) | 0.79 | 0.79 | 0.79 | 0.79 | 0.79 | 0.79 | 0.79 | 0.79 |
| $a$[9)] | 1.000 | 1.000 | 1.000 | 1.000 | 1.000 | 1.000 | 1.000 | 1.000 |
| $n_A$[10)] | 0.056 ± 0.001 | 0.049 ± 0.001 | 0.077 ± 0.001 | 0.067 ± 0.001 | 0.078 ± 0.004 | 0.066 ± 0.001 | 0.050 ± 0.001 | 0.051 ± 0.001 |
| $n_R$[11)] | 0.324 ± 0.001 | 0.278 ± 0.001 | 0.302 ± 0.002 | 0.204 ± 0.001 | 0.217 ± 0.004 | 0.234 ± 0.001 | 0.180 ± 0.001 | 0.187 ± 0.001 |
| $D$[12)] ($\mu m^2/s$) | 23.2 ± 0.1 | 25.7 ± 0.1 | 23.5 ± 0.1 | 24.8 ± 0.1 | 24.4 ± 0.1 | 27.2 ± 0.1 | 26.4 ± 0.1 | 27.8 ± 0.1 |
| $R_H$[13)] (nm) | 9.7 ± 0.0 | 8.3 ± 0.0 | 9.1 ± 0.0 | 8.6 ± 0.0 | 8.7 ± 0.0 | 7.8 ± 0.0 | 8.0 ± 0.0 | 7.6 ± 0.0 |

[1)]The ratio of the axial radius to the radial radius of the observation volume. [2)]The amplitude for the triplet state accumulation. [3)]The time constant attributed to the triplet state accumulation. [4)]The translational diffusion time for pCp-488. [5)]The translational diffusion time for 2′-OMe=tag-gRNA$_{12k}$-488. [6)]The brightness of the free Alexa488. [7)]The averaged fluorescence intensity. [8)]The background count rate. This value was determined 0.79 in our previous report (3). [9)]The ratio of the brightness of the labeled RNA relative to that of the free Alexa488. The value was fixed to 1 for the data in the absence of the N protein to estimate *b*, and set as a fitting parameter for the data in the presence of the N protein. [10)]The number of pCp-488 in the observation volume. [11)]The number of the labeled RNA in the observation volume. [12)]The diffusion coefficient and [13)]the hydrodynamic radius of 2′-OMe=tag-gRNA$_{12k}$-488.

**Supporting Table S3: The $R_H$ values of 2′-OMe=tag-gRNA$_{12k}$-488 at various concentration of NaCl and $MgCl_2$**

| [NaCl] (mM) | 0 | 0 | 0 | 0 | 150 | 150 | 150 | 150 |
|---|---|---|---|---|---|---|---|---|
| [$MgCl_2$] (mM) | 0 | 0.1 | 1 | 10 | 0 | 0.1 | 1 | 10 |
| $R_H$ (nm)_dataset1[1)] | 9.7 | 8.3 | 9.1 | 8.6 | 8.7 | 7.8 | 8.0 | 7.6 |
| $R_H$ (nm)_dataset2[1)] | 11.1 | 9.7 | 8.6 | 6.6 | 8.4 | 9.0 | 6.2 | 7.0 |
| $R_H$ (nm)_Average[2)] | 10.4 | 9.0 | 8.8 | 7.6 | 8.5 | 8.4 | 7.1 | 7.3 |
| $R_H$ (nm)_SD[3)] | 1.0 | 1.0 | 0.4 | 1.4 | 0.2 | 0.8 | 1.3 | 0.4 |

[1)]The hydrodynamic radius of 1.0–1.5 nM 2′-OMe=tag-gRNA$_{12k}$-488 estimated in the two independent measurements. [2)]The averaged $R_H$ for the 2 datasets. [3)]The standard deviation of $R_H$ for the 2 datasets.

**Supporting Table S4: The parameters used in and obtained by the fitting analysis of the representative FCS data set for 2 nM 2′-OMe=tag-gRNA$_{12k}$-488**

| [N] (nM) | 0 | 0 | 10 | 100 | 1000 |
|---|---|---|---|---|---|
| [NaCl] (mM) | 0 | 150 | 150 | 150 | 150 |
| [$MgCl_2$] (mM) | 0 | 1 | 1 | 1 | 1 |
| *s*[1] | 7.7648 | 7.7648 | 7.7648 | 7.7648 | 7.7648 |
| *frac*[2] | 0.185 ± 0.005 | 0.216 ± 0.004 | 0.211 ± 0.005 | 0.221 ± 0.008 | 0.265 ± 0.009 |
| $\tau_F$[3] (μs) | 3.7 ± 0.2 | 4.5 ± 0.2 | 4.3 ± 0.2 | 6.5 ± 0.4 | 5.6 ± 0.3 |
| $\tau_A$[4] (μs) | 107.2 | 107.2 | 107.2 | 107.2 | 107.2 |
| $\tau_R$[5] (μs) | 825.0 ± 2.6 | 825.0 ± 2.6 | 824.7 ± 2.8 | 1613.5 ± 10.9 | 2279.9 ± 19.0 |
| *B*[6] (photons/ms) | 14.88 ± 0.01 | 14.60 ± 0.02 | 14.60 | 14.60 | 14.60 |
| *I*[7] (photons/ms) | 9.9 | 8.1 | 6.4 | 2.0 | 2.1 |
| *BG*[8] (photons/ms) | 0.79 | 0.79 | 0.79 | 0.79 | 0.79 |
| *a*[9] | 1.000 | 1.000 | 0.929 ± 0.001 | 0.734 ± 0.001 | 0.648 ± 0.001 |
| $n_A$[10] | 0.090 ± 0.002 | 0.103 ± 0.002 | 0.075 ± 0.002 | 0.010 ± 0.001 | 0.012 ± 0.000 |
| $n_R$[11] | 0.522 ± 0.002 | 0.397 ± 0.002 | 0.333 ± 0.002 | 0.100 ± 0.001 | 0.120 ± 0.001 |
| *D*[12] (μm²/s) | 19.7 ± 0.1 | 27.1 ± 0.1 | 27.2 ± 0.1 | 13.9 ± 0.1 | 9.8 ± 0.1 |
| $R_H$[13] (nm) | 10.9 ± 0.1 | 7.8 ± 0.0 | 7.8 ± 0.0 | 15.2 ± 0.1 | 21.5 ± 0.2 |

[1]The ratio of the axial radius to the radial radius of the observation volume. [2]The amplitude for the triplet state accumulation. [3]The time constant attributed to the triplet state accumulation. [4]The translational diffusion time for pCp-488. [5]The translational diffusion time for 2′-OMe=tag-gRNA$_{12k}$-488. [6]The brightness of the free Alexa488. [7]The averaged fluorescence intensity. [8]The background count rate. This value was determined 0.79 in our previous report (3). [9]The ratio of the brightness of the labeled RNA relative to that of the free Alexa488. The value was fixed to 1 for the data in the absence of the N protein to estimate *b*, and set as a fitting parameter for the data in the presence of the N protein. [10]The number of pCp-488 in the observation volume. [11]The number of the labeled RNA in the observation volume. [12]The diffusion coefficient and [13]the hydrodynamic radius of 2′-OMe=tag-gRNA$_{12k}$-488.

**Supporting Table S5: The $R_H$ values of 2′-OMe=tag-gRNA$_{12k}$-488 at various concentrations of N protein**

| [N] (nM) | 0 | 0 | 10 | 100 | 1000 |
|---|---|---|---|---|---|
| [NaCl] (mM) | 0 | 150 | 150 | 150 | 150 |
| [$MgCl_2$] (mM) | 0 | 1 | 1 | 1 | 1 |
| $R_H$ (nm)_dataset1[1)] | 10.9 | 7.8 | 7.8 | 15.2 | 21.5 |
| $R_H$ (nm)_dataset2[1)] | 8.5 | 6.4 | 7.0 | 12.2 | 12.9 |
| $R_H$ (nm)_dataset3[1)] | 8.7 | 5.4 | 5.6 | 15.3 | 18.4 |
| $R_H$ (nm)_Average[2)] | 9.3 | 6.5 | 6.8 | 14.2 | 17.6 |
| $R_H$ (nm)_SD[3)] | 1.4 | 1.2 | 1.1 | 1.8 | 4.4 |

[1)]The hydrodynamic radius of 2′-OMe=tag-gRNA$_{12k}$-488 estimated in the three independent measurements. [2)]The averaged $R_H$ for the 3 datasets. [3)]The standard deviation of $R_H$ for the 3 datasets.

**Supporting Figures**

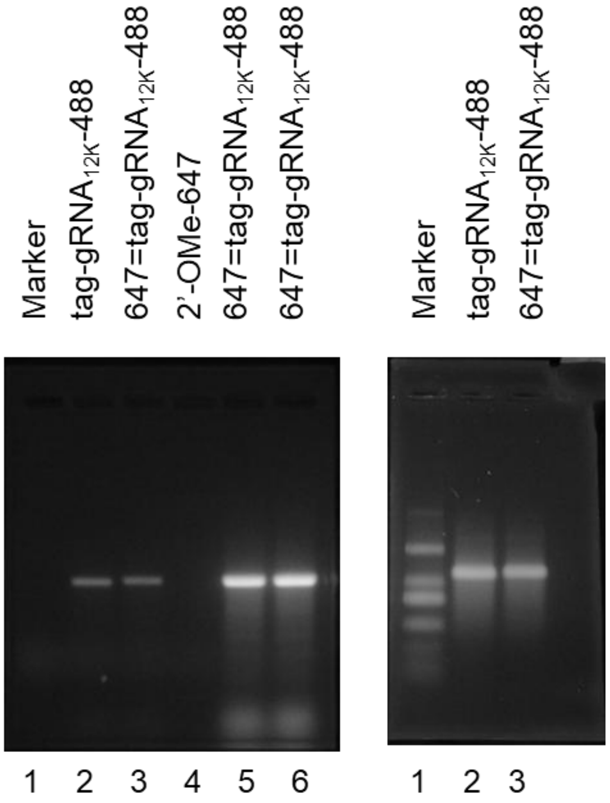


**Supporting Figure S1. Representative gel images for the labeling and purification steps of 647=tag-gRNA$_{12k}$–488**. After the annealing of 2′-OMe-647 with tag-gRNA$_{12k}$–488, the mixture was applied to the agarose gel for electrophoresis. The gel was irradiated with excitation light at 480–520 nm and imaged (left image). The intense signals in the 5th and 6th lanes were extracted and purified. The remaining gel including the 1st, 2nd, 3rd and 4th lanes was soaked with a SYBR gold solution and imaged to check the presence of non-labeled RNA samples (right image). The 1st, 2nd, 3rd and 4th lanes correspond to ssRNA maker, 0.13 μM tag-gRNA$_{12k}$–488, 0.13 μM 647=tag-gRNA$_{12k}$–488 and 0.13 μM 2′-OMe-647, respectively. The 5th and 6th lanes both correspond to 1.3 μM 647=tag-gRNA$_{12k}$–488.

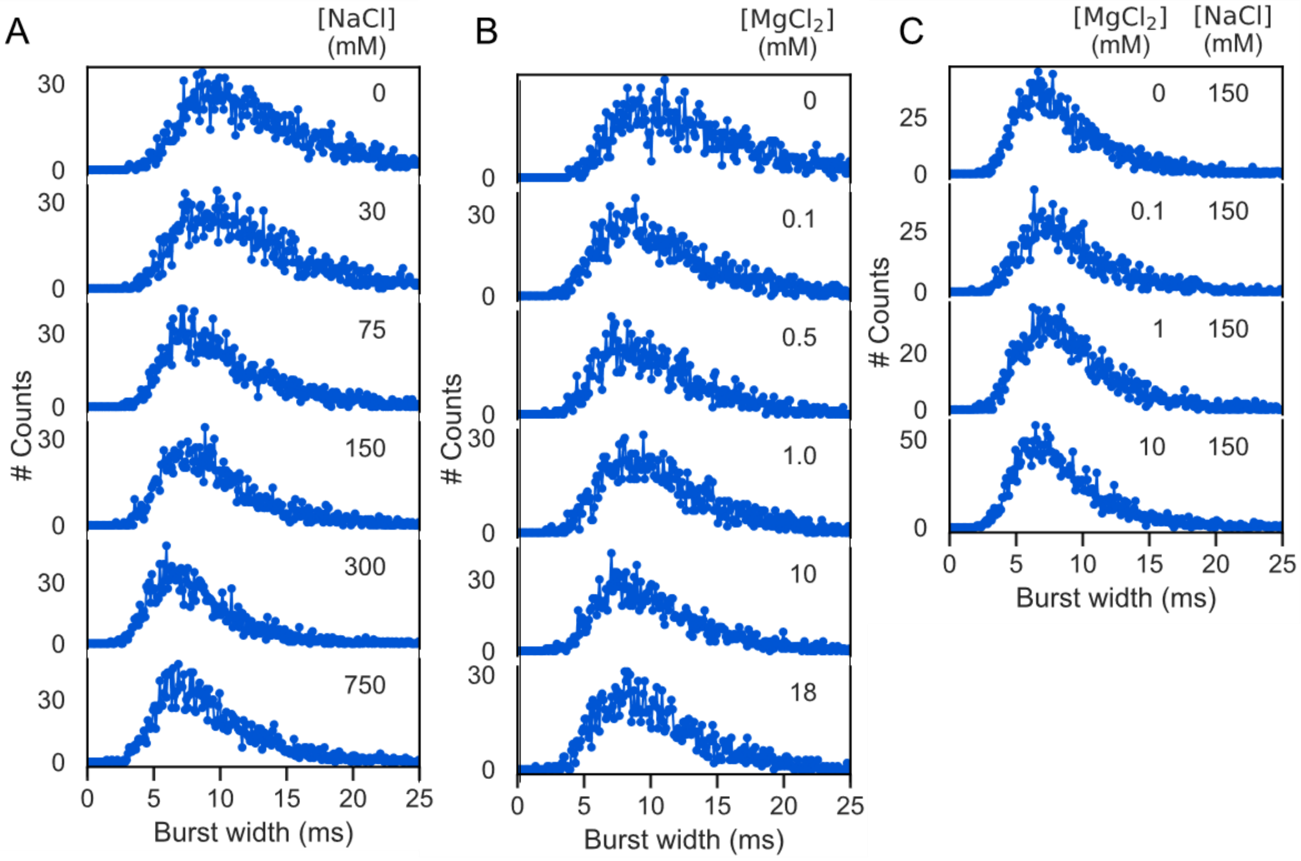


**Supporting Figure S2. The burst width distribution plots for the ALEX measurements of 50 pM 647=tag-gRNA$_{12k}$–488 in different concentrations of Na$^+$ and Mg$^{2+}$.** The horizonal axis and vertical axis are the burst width and the number of bursts, respectively. (A) The Na$^+$ concentration dependence in the absence of Mg$^{2+}$. (B) The Mg$^{2+}$ concentration dependence in the absence of Na$^+$. The gradual shortening of the burst width is consistent with the reduction of $R_H$ at the higher salt concentrations. (C) The Mg$^{2+}$ concentration dependence in the presence of 150 mM Na$^+$. All data were obtained without adding NaCl. The plots A, B and C correspond to the data presented as the FRET efficiency plots in Figs 2B, 2C and 2D, respectively, in the main text.

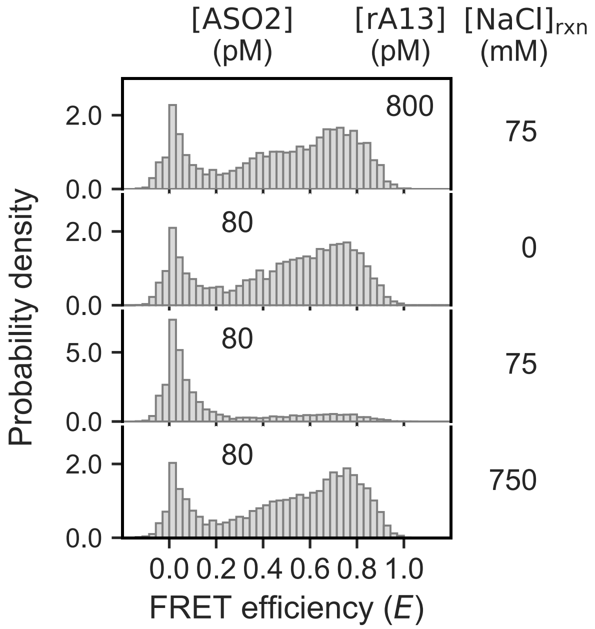


**Supporting Figure S3. The sm-FRET efficiency distribution for 647=tag-gRNA$_{12k}$–488 treated with ASO2 or rA$_{13}$ at various concentrations of NaCl.** The samples were prepared as explained in Supporting Text. [NaCl]rxn stands for the concentration of NaCl in the buffer during the hybridization reaction of ASO2 or rA$_{13}$ with 647=tag-gRNA$_{12k}$–488. In the absence of NaCl during the incubation in the presence of ASO2 at 100 nM, the replacement of the long-range base pairing with ASO2 did not occur (Second panel). Addition of 75 mM NaCl during the incubation promoted the hybridization of ASO2 (Third panel) but not that of rA$_{13}$ (Top panel). Addition of 750 mM NaCl during the incubation could not disrupt the high-efficiency peak even in the presence of ASO2 (Forth panel).

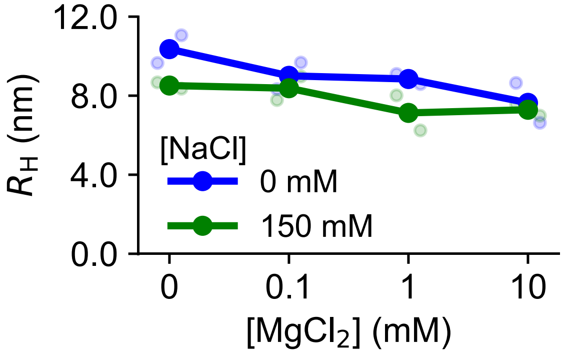


**Supporting Figure S4. $R_H$ of 2′-OMe=tag-gRNA$_{12k}$–488 at various concentrations of $Na^+$ and $Mg^{2+}$ determined by the analysis of the two independent FCS measurements excited at 488 nm.** Blue and green dots represent the mean $R_H$ values of the two independent measurements obtained in the absence and presence of 150 mM $Na^+$, respectively. Light dots indicate the result of individual measurements. The detailed fitting parameters are presented in Supporting Table S2. The averaged $R_H$ values are presented in Supporting Table S3.

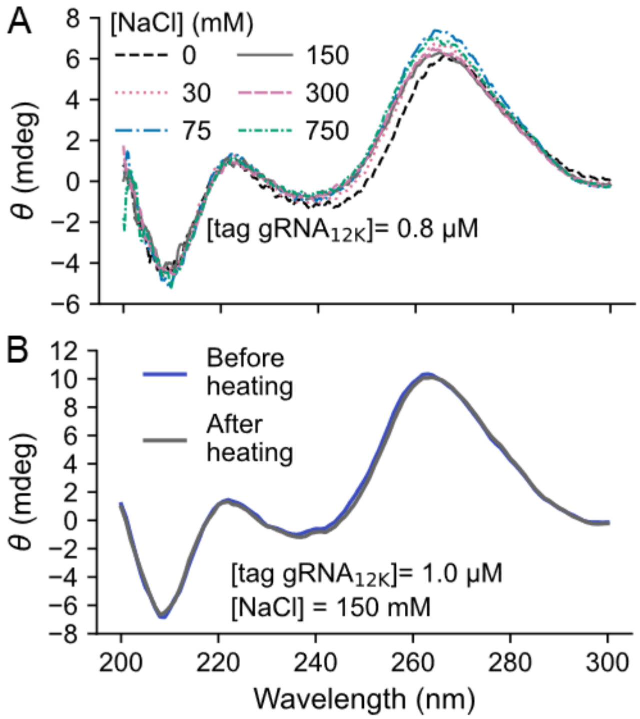


**Supporting Figure S5. CD spectra of tag-gRNA$_{12k}$.** (A) CD spectra obtained at room temperature for 0.8 µM tag-gRNA$_{12k}$ at the different concentrations of NaCl. (B) CD spectra of 1.0 µM tag-gRNA$_{12k}$. The spectra were measured at room temperature before and after the heat treatment at 85°C.

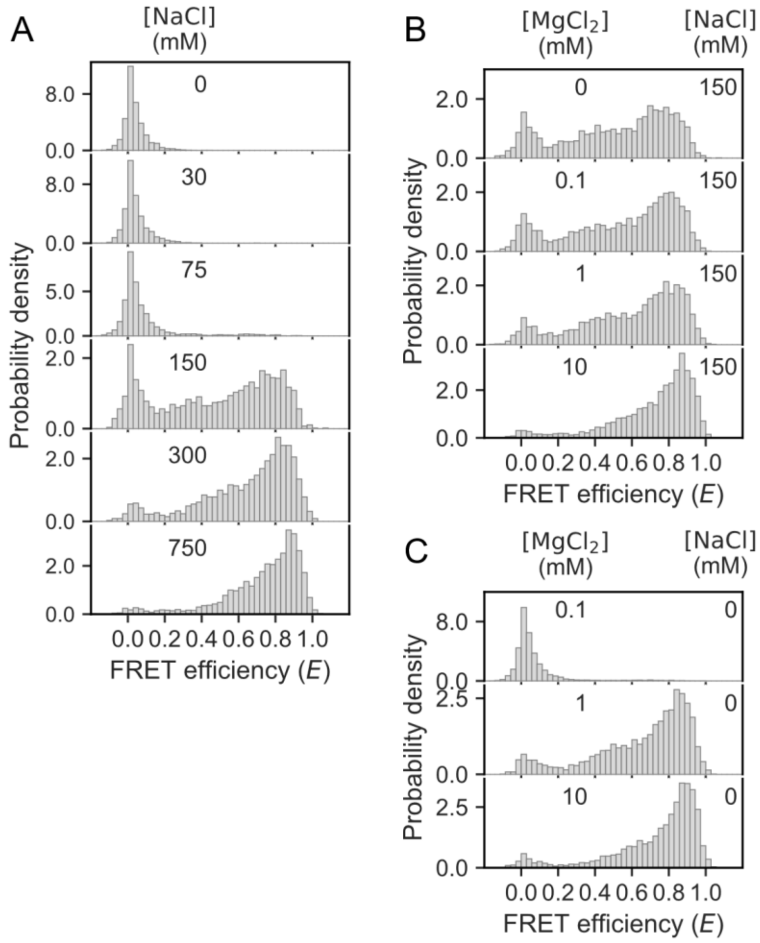


**Supporting Figure S6. The sm-FRET efficiency distribution for 647=tag-gRNA$_{12k}$–488 incubated at 75°C for 5 min followed by the mixing with varying concentrations of cations at room temperature.** The FRET efficiency plots were obtained at the varying concentration of $Na^+$ (A), at the varying concentrations of $Mg^{2+}$ in the presence of 150 mM $Na^+$ (B), and at the varying concentration of $Mg^{2+}$ in the absence of $Na^+$ (C). The concentration of the labeled RNA was 50 pM. The concentrations of NaCl and $MgCl_2$ are described in each panel. More than 1900 bursts were collected to depict distributions in each panel.

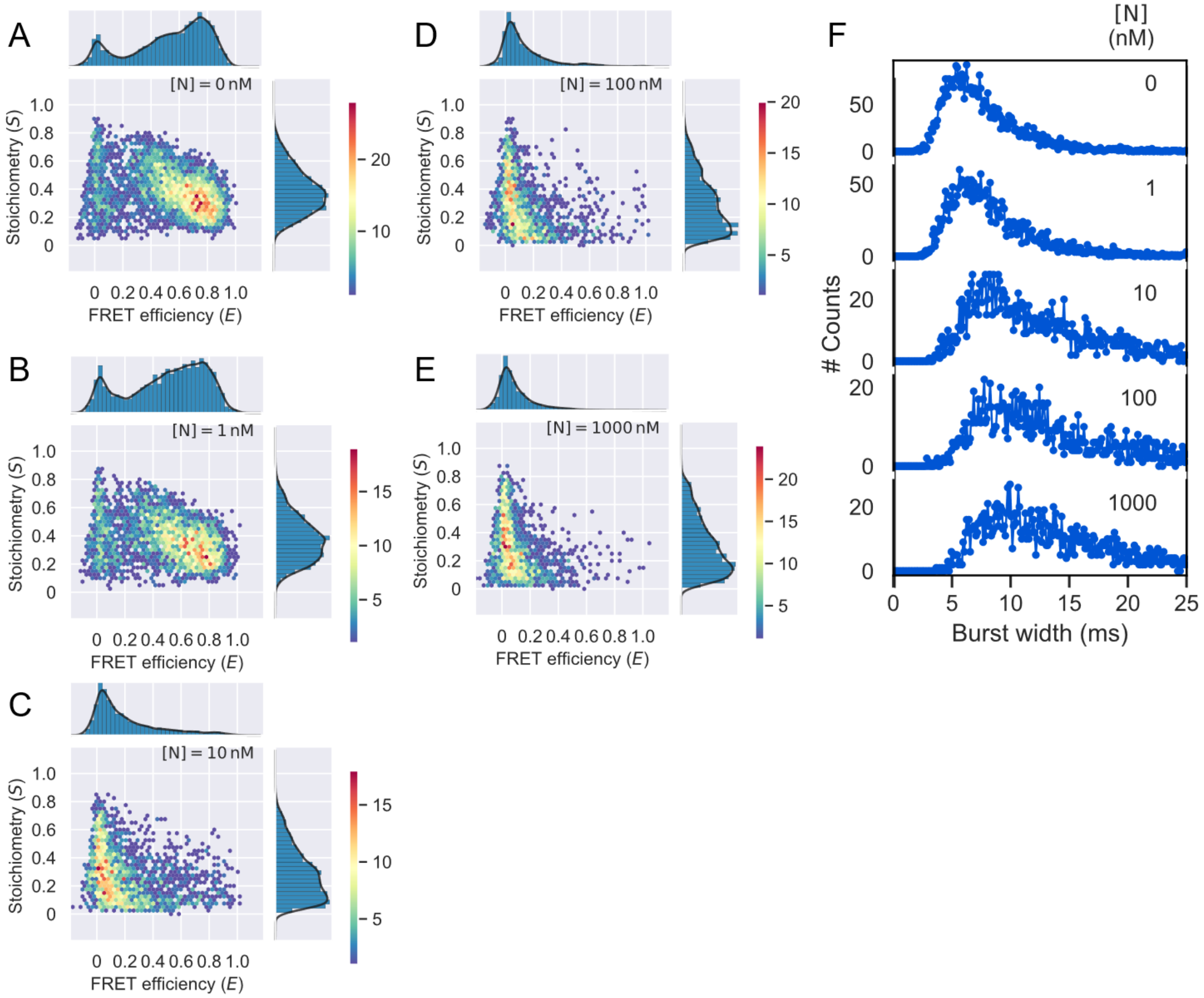


**Supporting Figure S7. The two-dimensional *E*-*S* plots (A–E) and the burst width plots (F) for 647=tag-gRNA$_{12k}$–488 obtained at varying concentrations of the N protein and 150 mM Na$^+$.** (A–E) The horizonal and vertical axes are FRET efficiency (*E*) and stoichiometry (*S*), respectively. Color bars indicate the numbers of bursts. (F) The horizonal axis and vertical axis are the burst width and the number of bursts, respectively. These plots correspond to Fig. 5A in the main text. The concentrations of the labeled RNA and NaCl were 100 pM and 150 mM, respectively. The concentrations of the N protein are described in each panel. More than 1800 bursts were collected to depict distributions in each panel.

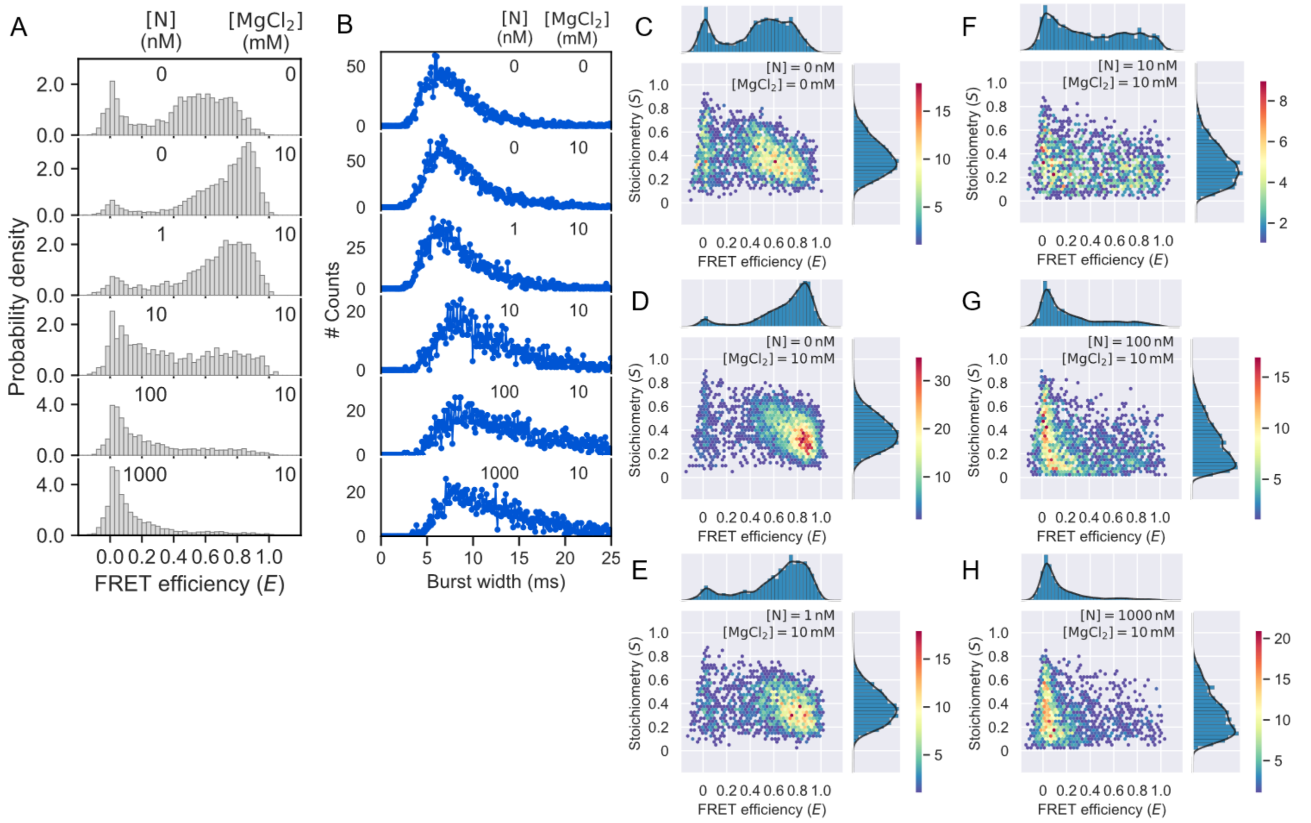


**Supporting Figure S8. The sm-FRET measurements for 647=tag-gRNA$_{12k}$–488 obtained at varying concentrations of the N protein in the presence of 10 mM $Mg^{2+}$ and 150 mM $Na^{+}$.** (A) Changes in the sm-FRET efficiency distribution. (B) Changes in the burst width distribution. (C–H) The two-dimensional scattering plots whose horizonal and vertical axes represent the sm-FRET efficiency and stoichiometry, respectively. Color bars indicate the numbers of bursts. For all the data, the concentrations of the labeled RNA and NaCl were 100 pM and 150 mM, respectively. More than 1500 bursts were collected to depict distributions in each panel.

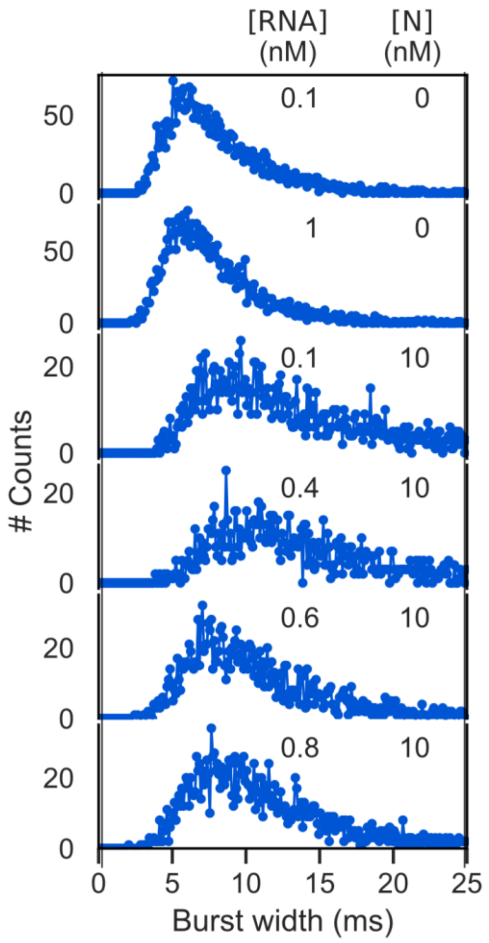


**Supporting Figure S9. The burst width distribution plots in the ALEX measurements of 647=tag-gRNA$_{12k}$ at varying concentrations of the N protein and the non-labeled tag-gRNA$_{12k}$.** These plots correspond to Figure 5D in the main text. The concentrations of the labeled RNA and NaCl were 100 pM and 150 mM, respectively. The concentrations of the N protein are described in each panel. The total concentration of 647=tag-gRNA$_{12k}$–488 and non-labeled tag-gRNA$_{12k}$ are described in each panel as [RNA].

### Supporting References


1. K. Kuwahara, S. Yajima, Y. Yamano, F. Nagatsugi, K. Onizuka, Formation of Direction-Controllable Pseudorotaxane and Catenane Using Chemically Cyclized Oligodeoxynucleotides and Their Noncovalent RNA Labeling. *Bioconjugate Chem.* (2023).

2. G. J. Smith, T. R. Sosnick, N. F. Scherer, T. Pan, Efficient fluorescence labeling of a large RNA through oligonucleotide hybridization. *RNA* **11**, 234–239 (2005).

3. N. Kaneda, *et al.*, A Single Dimer of the SARS-CoV-2 N Protein Can Associate with Multiple Fragments of Single-Stranded and Stem-Loop RNA: A Single-Molecule FRET and FCS Investigation. *ACS Omega* **11**, 21382–21394 (2026).

4. A. Ingargiola, T. Laurence, R. Boutelle, S. Weiss, X. Michalet, Photon-HDF5: An Open File Format for Timestamp-Based Single-Molecule Fluorescence Experiments. *Biophysical Journal* **110**, 26–33 (2016).

5. B. Hellenkamp, *et al.*, Precision and accuracy of single-molecule FRET measurements-a multi-laboratory benchmark study. *Nat Methods* **15**, 669–676 (2018).

6. A. Ingargiola, E. Lerner, S. Chung, S. Weiss, X. Michalet, FRETBursts: An Open Source Toolkit for Analysis of Freely-Diffusing Single-Molecule FRET. *PLoS ONE* **11**, e0160716 (2016).

7. S. Mitra, *et al.*, Flexible Target Recognition of the Intrinsically Disordered DNA-Binding Domain of CytR Monitored by Single-Molecule Fluorescence Spectroscopy. *J. Phys. Chem. B* **126**, 6136–6147 (2022).

8. J. Kestin, H. E. Khalifa, R. J. Correia, Tables of the dynamic and kinematic viscosity of aqueous NaCl solutions in the temperature range 20–150 °C and the pressure range 0.1–35 MPa. *Journal of Physical and Chemical Reference Data* **10**, 71–88 (1981).

9. P.-O. Gendron, F. Avaltroni, K. J. Wilkinson, Diffusion Coefficients of Several Rhodamine Derivatives as Determined by Pulsed Field Gradient–Nuclear Magnetic Resonance and Fluorescence Correlation Spectroscopy. *J Fluoresc* **18**, 1093–1101 (2008).

10. M. A. Digman, R. Dalal, A. F. Horwitz, E. Gratton, Mapping the Number of Molecules and Brightness in the Laser Scanning Microscope. *Biophysical Journal* **94**, 2320–2332 (2008).

11. Y. Chen, J. D. Müller, P. T. C. So, E. Gratton, The Photon Counting Histogram in Fluorescence Fluctuation Spectroscopy. *Biophysical Journal* **77**, 553–567 (1999).

12. H. Qian, E. L. Elson, Distribution of molecular aggregation by analysis of fluctuation moments. *Proceedings of the National Academy of Sciences* **87**, 5479–5483 (1990).
